# An approximate full-likelihood method for inferring selection and allele frequency trajectories from DNA sequence data

**DOI:** 10.1101/592675

**Authors:** Aaron J. Stern, Peter R. Wilton, Rasmus Nielsen

**Affiliations:** Graduate Group in Computation Biology, University of California, Berkeley, Berkeley, CA 94702; Department of Integrative Biology, University of California, Berkeley, Berkeley, CA 94702; Department of Statistics, University of California, Berkeley, Berkeley, CA 94702

## Abstract

Most current methods for detecting natural selection from DNA sequence data are limited in that they are either based on summary statistics or a composite likelihood, and as a consequence, do not make full use of the information available in DNA sequence data. We here present a new importance sampling approach for approximating the full likelihood function for the selection coefficient. The method treats the ancestral recombination graph (ARG) as a latent variable that is integrated out using previously published Markov Chain Monte Carlo (MCMC) methods. The method can be used for detecting selection, estimating selection coefficients, testing models of changes in the strength of selection, estimating the time of the start of a selective sweep, and for inferring the allele frequency trajectory of a selected or neutral allele. We perform extensive simulations to evaluate the method and show that it uniformly improves power to detect selection compared to current popular methods such as nSL and SDS, under various demographic models and can provide reliable inferences of allele frequency trajectories under many conditions. We also explore the potential of our method to detect extremely recent changes in the strength of selection. We use the method to infer the past allele frequency trajectory for a lactase persistence SNP (*MCM6*) in Europeans. We also study a set of 11 pigmentation-associated variants. Several genes show evidence of strong selection particularly within the last 5,000 years, including *ASIP*, *KITLG*, and *TYR*. However, selection on *OCA2/HERC2* seems to be much older and, in contrast to previous claims, we find no evidence of selection on *TYRP1*.

**Author summary:** Current methods to study natural selection using modern population genomic data are limited in their power and flexibility. Here, we present a new method to infer natural selection that builds on recent methodological advances in estimating genome-wide genealogies. By using importance sampling we are able to efficiently estimate the likelihood function of the selection coefficient. We show our method improves power to test for selection over competing methods across a diverse range of scenarios, and also accurately infers the selection coefficient. We also demonstrate a novel capability of our model, using it to infer the allele’s frequency over time. We validate these results with a study of a lactase persistence SNP in Europeans, and also study a set of 11 pigmentation-associated variants.

## Introduction

Direct observation of the change in allele frequency over time (the allele frequency trajectory) allows one to make powerful inferences regarding whether selection acted on the allele [1,2]. However, outside of certain contexts such as experimental evolution of viruses or bacteria [3–6] or analyses of ancient DNA samples [7,8], in most cases such direct observations of allele frequencies at multiple points in the history of a population are unavailable. Instead, selection must be inferred from contemporary, modern data. A wide variety of methods have been developed to detect selection based on patterns observed from modern DNA sequences (e.g. [9–11]).

The hitch-hiking effect provides a key signature of selection in modern datasets [12,13]. Hitch-hiking causes aberrations in the spatial pattern of genetic diversity, including the site frequency spectrum (SFS) [14,15] and the pattern of haplotype homozygosity [9]. Methods designed to detect these aberrations are particularly useful in the setting where a single population is surveyed, and the only information available is variation within this single population.

The most familiar methods for detecting selection are based on linear functions of the SFS, such as Tajima’s *D*, Fu and Li’s *D*, or Fay and Wu’s *H* [14,16,17]. An advantage of SFS-based methods is that they do not require the data to be phased. However, these methods have several limitations: they tend to confound selection with other non-equilibrium conditions, such as a fluctuating population size [10,18]; they are not suitable for estimating parameters such as the value of the selection coefficient *s*; significance can usually only be established using an empirical null distribution; and crucially, these methods do not incorporate any features of the haplotype structure.

To make fuller use of information provided by phased sequence data, a number of methods have incorporated summary statistics based on haplotype structure. In a broad sense, these methods are based on calculations of haplotype similarity in a window around some core site of interest [9]. Several methods have adapted this general concept to specifically detect ongoing selection [11,19,20]. More recently, [21] showed that the density of singletons surrounding a focal SNP can be a powerful signal of extremely recent selection in large cohorts. In addition to recent and ongoing selection, it has been demonstrated that these methods have compelling advantages to detecting selection from standing variation [20–22]. However, these methods share the major limitation of SFS-based method in that they are not suitable for parametric inference and it is unclear how to establish significance without use of an empirical null model.

Recently, supervised machine learning methods have been proposed as an alternative to traditional summary-statistic based methods (see e.g., [23]). Standard machine learning techniques applied to population genomic data afford some major advantages over methods based on summary-statistics: standard techniques can produce accurate classifiers based on summaries of the data that live in much higher-dimensional space than the aforementioned summary statistics, and these techniques often encompass a wide space of classification functions that are often non-linear (see e.g. [24,25]). Some studies have demonstrated these methods can have improved robustness to demographic model mis-specification [22,26]. Although these methods can potentially detect complex patterns left by selection, they demand training on large data sets which typically are simulated using models that may not accurately correspond to the empirical data.

In contrast to the aforementioned methods, one might aim to develop a full likelihood method which would take into account the full data set, rather than merely summary statistics. A common strategy for obtaining the full likelihood has been to find the distribution of the genealogy under selection. For example, Krone and Neuhauser described the distribution of the coalescence tree of a locus under weak selection and no recombination [27]. Alternatively, one can describe how the genealogy depends on the trajectory of the derived allele (first described by [28]), and in turn how the trajectory depends on selection. To this end, Coop and Griffiths [29] developed a sampling method for approximating the full likelihood of the selection coefficient. Their method uses sampling to marginalize out two layers of latent variables: the allele frequency trajectory and the genealogy of the locus. To estimate the likelihood function, they perform random sampling of both the trajectory, and the genealogy conditioned on the trajectory. Unfortunately, selection likelihood methods that consider the both coalescence and recombination are generally considered computationally intractable.

Composite likelihood methods (see e.g. [10,30]) are able to approximate the likelihood function using tractable expressions for the frequency distribution of a neutral site linked to the selected site [15,31]. These methods approximate the joint distribution of frequencies observed at linked sites as the product of their marginals. These approaches can be applied to test for selection, and estimate the strength of selection. The approximations made by composite likelihood methods are more accurate under strong selection (arguably beyond the strength of most recent selection in humans), and thus have less power to detect weak selection — although to some extent low power to detect weak selection is a natural outcome of any selection method.

Approximate Bayesian computation (ABC) and rejection sampling methods approximate the likelihood function by simulation. One advantage over the composite likelihood approach is that ABC can capture dependencies between linked neutral sites. For example, methods have been used to jointly infer the strength and timing of selection acting on a locus and determine whether a sweep occurred from a *de novo vs* standing variant [32–35]. However, a major disadvantage of such approaches is that the amount of simulation necessary to obtain an accurate estimate grows dramatically with the dimensionality of the model parameters. There is an additional tradeoff between information utilized from the data and computational burden; as the number of the summary statistics used increases, the number of simulations required to approximate the likelihood at a fixed parameter value also increases (for a discussion, see e.g. [36]).

The method we present in this paper draws inspiration from the Coop & Griffiths method [29], and has several key similarities: our method produces a likelihood; involves integrating out the allele frequency trajectory and genealogy, i.e., the aforementioned two hidden layers; and both methods account for selection by modeling how allele frequency changes depend on selection. However, there are several key differences between this method and our approach: while Coop and Griffiths assume no recombination of the locus, our method is based on the coalescent with recombination (i.e. the ancestral recombination graph or ARG) [37]. Also, whereas Coop & Griffiths simulate random trajectories, we use dynamic programming algorithms similar to those used in Hidden Markov Models (HMMs) to completely marginalize the latent trajectory. The hidden states represent allele frequencies and the emission probabilities are coalescence probabilities. While the framework is not a traditional HMM because the process is time-inhomogenous and the emission space changes with time, the similarities with traditional HMMs are, nonetheless, so significant that we will refer to this as an HMM. Lastly, our method uses a novel importance sampling scheme that allows us to sample ARGs assuming a neutral prior, and find the likelihood function at arbitrary values of *s*; this drastically reduces the amount of ARG sampling necessary.

Furthermore, the new method is, to our knowledge, the first that is capable of inferring the allele frequency trajectories for models with recombination and selection using only modern data. We are able to accomplish this task using the aforementioned Markovian structure of both coalescence and the trajectory, forming a HMM over these two hidden states and solving for the posterior marginals of each hidden allele frequency state over time. Recently, Edge & Coop proposed a method to reconstruct changes to polygenic scores over time via such estimates of the local trees, but their method is not suitable for estimating allele frequency changes or selection at individual loci [38].

## Materials and methods

### Overview

We begin with an overview of our method for jointly inferring selection and the allele frequency trajectory, which we summarize in Fig. 1. Our method begins with input in the form of phased SNP data from a linked genomic region (Fig. 1A), although technically, it is also possible to use unphased data, and sample possible phasings. While the method generalizes to arbitrary sample size, we recommend using *n* = 25 *−* 100 diploid individuals when using ARGweaver as done in this study, as ARGweaver runtime increases quadratically with sample size. The method also generalizes to arbitrarily long regions, although we recommend using regions of 10^2^ *−* 10^3^kb, roughly the size of many LD blocks in the human genome [39].

**Fig 1.**
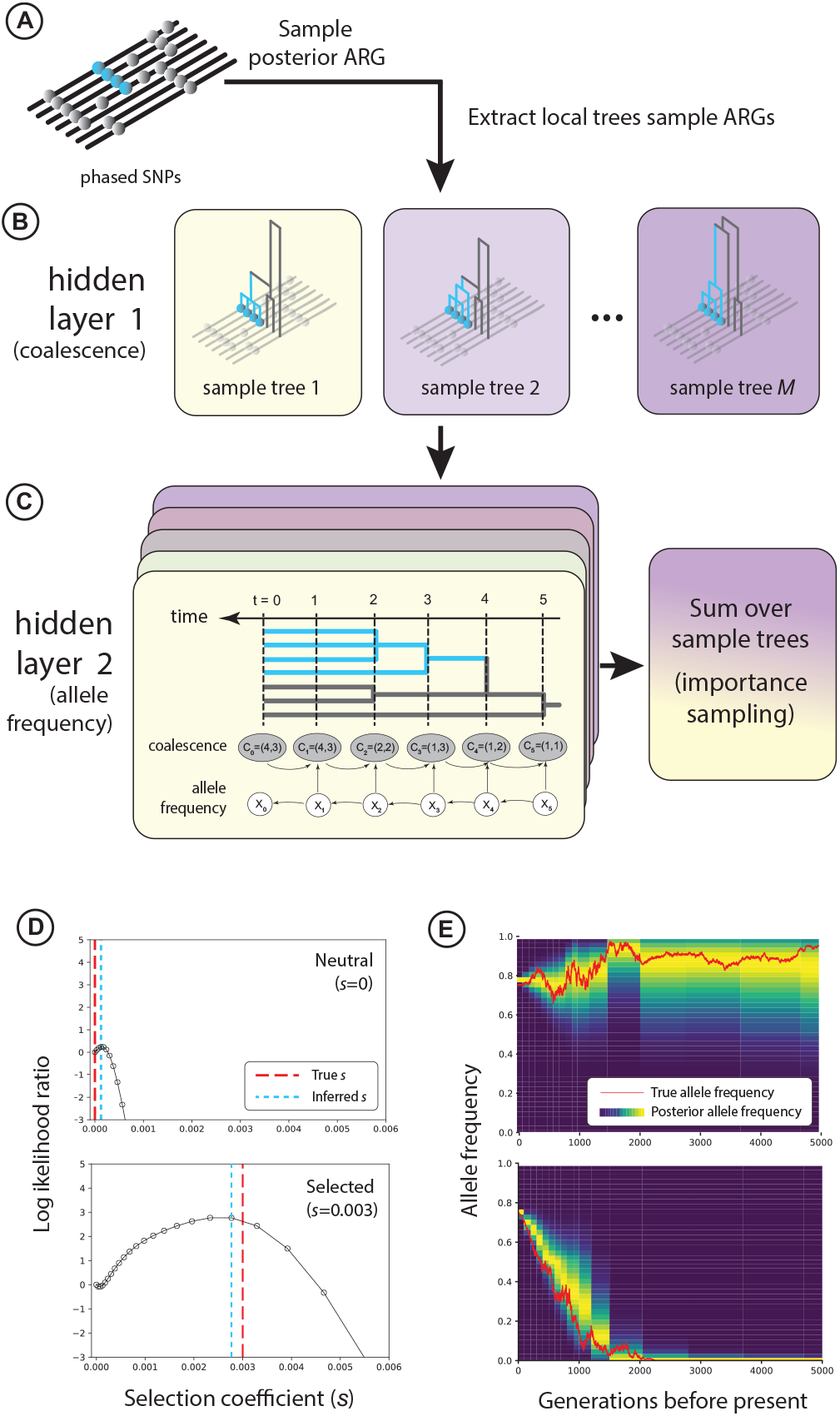
A: To apply our method for inferring selection, we begin by sampling the posterior ARG of a set of recombining chromosomes. B: For each sample ARG, we extract local trees at the site of interest (blue). C: For each sample local tree, we run an HMM to calculate the likelihood of selection, marginalizing out the hidden allele frequency trajectory based on coalescence in the sample tree. We later use the recursions performed in this step to calculate the posterior allele frequency trajectory. D: An example of the estimated likelihood function for an allele under neutrality (top) and selection (bottom). E: An example of the inferred allele frequency trajectory compared to the ground truth trajectory under neutrality (top) and selection (bottom). Both (D) and (E) are inferred from data simulated under a European demographic model with *n* = 50 haplotypes, conditioning on the derived allele segregating at 75% in the present day. with *s* = 0 and *s* = 0.003, respectively.

Next, we sample the genealogy at the selected site from its posterior distribution, assuming a neutral model (Fig. 1B). By sampling over this distribution, we marginalize out the hidden coalescence events, the first of two latent variables or “hidden layers” in our model. Specifically, we sample the full ancestral recombination graph (ARG) of the input haplotypes. The ARG is a graph that summarizes all of the common ancestry and recombination events that have occurred within the sample. We sample ARGs rather than gene trees in order to account for recombination, and to incorporate information from sites in long-range linkage disequilibrium with the selected site. Then we extract the genealogy at the site of interest (the “local tree”) and from here on, this is the only component of the ARG that goes into our subsequent calculations. To perform ARG sampling, we choose to use ARGweaver [37], which is the only currently available method to sample the posterior ARG. In practice, it is possible and straightforward to adapt this method to other ARG inference methods designed for larger samples, but sampling the posterior yields beneficial statistical properties (see “Importance Sampling” under Materials and Methods).

Then, for each local tree we have sampled, we form a hidden Markov model (HMM, Fig. 1C) indexed in time according to the discretization chosen for ARGweaver; in this HMM, observed states are coalescence times in this local tree and hidden states are the selected allele’s frequency trajectory over time (i.e., the second hidden layer of our overall model). We use a discrete-time model of the coalescent process to match the model used by ARGweaver, so that the length of the HMM is of manageable, finite length. Emission probabilities (i.e., coalescence probabilities) depend both on the allele frequency and the current coalescent state. Hence, the model is time-inhomogenous as the coalescent state changes through time. However, the dependence structure is otherwise identical to traditional HMMs and all the usual dynamic programming algorithms apply. The transition probabilities of allele frequencies depend on the selection coefficient *s*, the parameter we are ultimately interested in estimating. Marginalizing out the allele frequency trajectory from the HMM yields the probability of the sample local tree as a function of *s*. To obtain the likelihood function of *s*, we perform importance sampling over all sample trees, reweighting their coalescent probabilities and summing them up. This approach allows us to use trees sampled exclusively under a prior of selective neutrality (*s* = 0) to calculate the likelihood function at arbitrary values of *s*. In other words, this approach allows us to minimize the amount of ARG sampling necessary to estimate the likelihood function, which is notable because ARG sampling is generally the most computationally intensive step of our method.

Finally, we can analyze the results to test for selection or estimate the selection coefficient (Fig. 1D). Additionally, we show that we can decode the HMMs depicted in Fig. 1C and use them to obtain a posterior estimate of the allele frequency trajectory (Fig. 1E).

A glossary to accompany the following derivations and description of the method is available in S1 Text.

### Coalescent model for a site under selection

First, let us consider how the distribution of the local tree *T* at a site under selection depends on the frequency trajectory of an allele at that site. We assume that the tree is labeled, i.e. we know which branches subtend each allele. We also assume the tree to be compatible with the infinite sites assumption, i.e. that there is at most one mutation event that has occurred at the focal site, and thus the site is bi-allelic. We model the likelihood of the tree using a structured coalescent; moving backwards in time from the time of sampling until the time of the mutation, lineages can only coalesce with other lineages that subtend the same allele, and the coalescence rate within the derived and ancestral classes depends on both the derived allele frequency *X*(*t*) and the effective population size *N* (*t*), both indexed by the time *t ≥* 0 in coalescent units before the present day. Proceeding back in time, lineages coalesce freely after the time of mutation, and the coalescence rate depends only on *N* (*t*). In the rest of this section we treat the trajectory *X*(*t*) as known, but in practice the trajectory is hidden and highly stochastic; in a later section we develop a hidden Markov model to efficiently integrate out *X*(*t*).

We use a discrete-time model of the coalescent employed also by ARGweaver [37]. That is, we only observe the coalescent process at a set of *K* discrete timepoints {*t*_1_,…, *t*_*K*_}, and also make the additional assumption that all lineages must coalesce by *t*_*K*_. (Typically *t*_*K*_ is set to *∼*100× *N*_*e*_, implying coalescence would be extremely unlikely to occur after *t*_*K*_, and hence this assumption is very reasonable.) Henceforth, using this discretization we also discretize *X* and *N*; we assume *X*(*t*) = *X*_*i*_ for *t* ∈ (*t*_*i*_, *t*_*i*+1_], and *N* (*t*) = *N*_*i*_ for *t* ∈ (*t*_*i*_, *t*_*i*+1_].

We use *C* to track the number of lineages remaining at these timepoints leading back into the past; as long as we keep track of the number of lineages belonging to each of the allelic classes, by exchangeability of lineages within an allelic class, we can model the likelihood function in the usual way, as independent of the topology given the waiting times. Hence, we define three simultaneous, related processes *C* = (*C*^der^, *C*^anc^, *C*^mix^). The processes *C*^der^ and *C*^anc^ refer to coalescence within the derived and ancestral classes during the time going back from the time of sampling to the time of the mutation. The mixed process *C*^mix^ refers to coalescence going backwards from the time of the mutation. We call it the mixed process because it includes un-coalesced lineages from *C*^anc^, as well as the lineage ancestral to all derived lineages. Assuming the infinite sites model, *C*^mix^ will have one additional lineage relative to *C*^anc^ at the time of the mutation, and will eventually reach *C*^anc^ = *C*^mix^ once that lineage coalesces with one of the other lineages in the ancestral class. In Fig. 2 we illustrate the lines-of-descent process in the these three classes.

**Fig 2.**
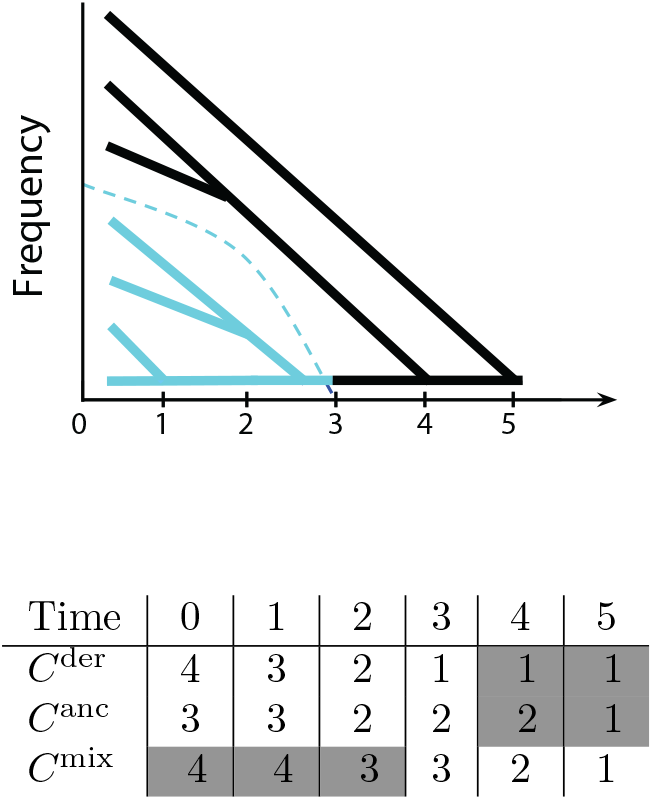
Top: Coalescence conditioned on the allele frequency trajectory (dashed blue line). Blue lineages subtend the derived allele, whereas black lineages do not. Black lineages belong to the ancestral class while the derived allele has *X*_*t*_ > 0, and they belong to the mixed class while *X*_*t*_ = 0. Bottom: the numbers of derived, ancestral, and mixed lineages at each time point. Numbers with unshaded cells factor into the likelihood calculation, whereas numbers with shaded cells do not.

We model the probability of transitioning from *C*_*i*_ → *C*_*i*+1_ lineages during some time interval [*t*_*i*_, *t*_*i*+1_] using a simple variation of Tavare’s formula for the exact distribution of the number of lines of descent remaining after *t* generations [40]. We use Tavare’s formula in order to model the coalescent at discrete timepoints, allowing multiple coalescences at each epoch.

We write the probability of *C* given the trajectory *X* (note this is distinct from the full likelihood of *s*) as

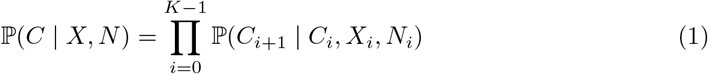

More precisely, in terms of the derived, ancestral, and mixed processes,

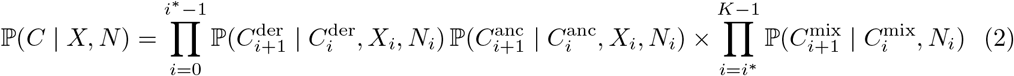

where *i*^∗^:= max{*i*: *X*_*i*_ > 0} denotes the index of the epoch during which the allele arose via mutation. Naturally, the mixed process—which we only keep track of while the derived allele is nonexistent—does not depend on *X*. We can write the transition probabilities using a variation of Tavare’s formula for the transition probabilities of the number of lines of descent [40]; in place of the effective population size, we substitute the size of a allelic class *z*^class^ in order to reflect the coalescence rate within an allelic class:

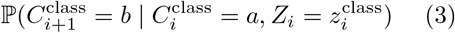

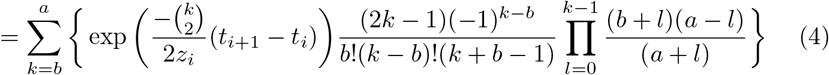

where

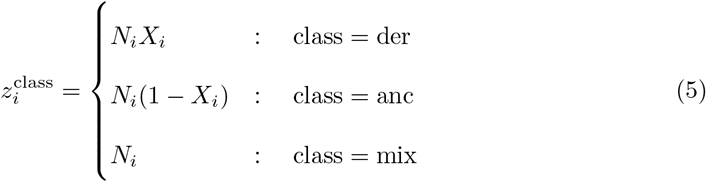

We note that this formula is known to be computationally unstable for large values of *C*, large values of *N*, and/or small values of ∆*t*_*i*_ = *t*_*i*+1_ − *t*_*i*_; under such conditions, the asymptotic distribution of 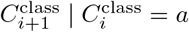(where *a* is, e.g., the number of derived lines of descent present at *t*_*i*_) takes on a normal distribution [41]:

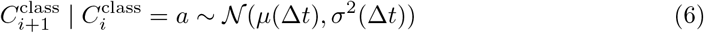

where

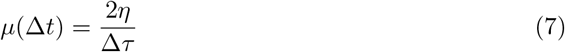

and

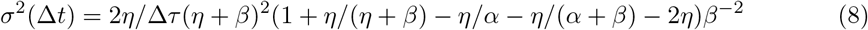

and

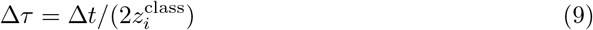

where *α* = *a*∆*τ*/2, *β* = *−*∆*τ*/2, and *η* = *αβ*/[*α*(*e*^*β*^ − 1) + *βe*^*b*^] [41]. In practice, for samples of *n* = 50 haplotypes under constant *N*_*e*_ = 10^4^, we find this approximation is unnecessary; however, for the same sample size under a European demographic model, which exhibits very large recent *N*_*e*_, we find it necessary to use this approximation during the roughly 10^3^ generations preceding the present day, prior to which ∆*t* is sufficiently large that we change over to Tavare’s exact formula [42].

### Marginalizing the hidden allele frequency states

In the previous sections we showed how we obtain 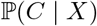 and 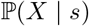. Here we illustrate how to model coalescence in the two allelic classes when the trajectory *X* is as a random latent variable. The probability of *C* given *s* is thus

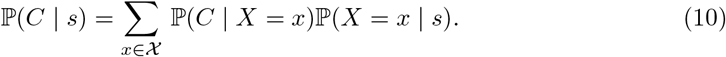

Naively, this involves a prohibitively large sum over *d*^*K*−1^ terms in 𝓧, the space of possible trajectories. But due to the conditional independence of the likelihood, we can calculate the likelihood much faster using a recursion similar to the forward and backward algorithms commonly deployed on HMMs; we show the derivations of these recursions in S1 Text. In essence, this algorithm works by iteratively marginalizing out the allele frequency in each epoch; the backward recursion marginalizes frequency during earlier epochs first and later epochs last, and the forward recursions marginalizes in reverse order. At each epoch *i* the recursions yield *b*_*i*_(*x*_*i*_) and *f*_*i*_(*x*_*i*_); these correspond to 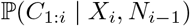 and 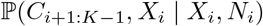, where *C*_*a:b*_ = *C*_*a*_, *C*_*a*+1_,…, *C*_*b*_.

Consequently, the two recursions can be used together to obtain the the posterior probability of the allele frequency during the *i*th epoch *X*_*i*_,

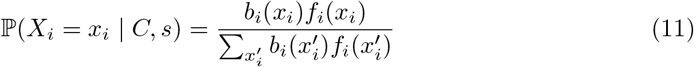

which gives the posterior marginal of *X*_*i*_ using the familiar forward-backward algorithm.

### Importance sampling to estimate the likelihood function

The above formulas pertain immediately only to the case in which the local tree is observed directly and without noise. In practical settings, the local tree is hidden to us and we must integrate over the space of possible local trees using sampling methods. Here we describe a novel importance sampling method to reweight posterior samples of the ARG to approximate the likelihood function of selection. Although we use *s* to express the argument of the likelihood function, we use this as shorthand for estimating the likelihood function of arbitrarily complex parameters; for example, one could estimate the selection coefficient *s*, as well as the time of selection’s onset, *t*_*s*_, before which the allele behaved neutrally.

We are given haplotype data *D* representing *n* haplotypes with *l* sites that are fixed for the derived allele. We wish to use *D* to infer the maximum-likelihood value of *s* for some site *k* ∈ {1,2,…, *l*}, where *l* is the number of sites in the region, assuming that all other sites are selectively neutral (i.e. *s*_*j*_ = 0 ∀*j* ∈ {1, 2,…, *k* − 1, *k* + 1,…, *l*}). In other words, we restrict ourselves to testing simple hypotheses of the form “site *k* has selection coefficient *s*_*k*_ and all of its flanking sites are selectively neutral.”

The likelihood of *s* under the data can be expressed as the expected value of the likelihood of the ARG 𝓖 given the data *D*, with respect to the distribution of 𝓖 given *s*:

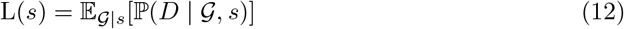

At this stage, we introduce *G*, the discrete-time approximation of 𝓖 (discussed in more detail by [37]), and we assume

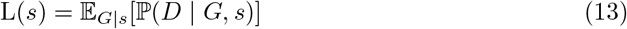

By importance sampling, we are able to express the expectation over an alternative distribution *q*(*G*), as long as 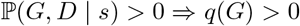. Notice that this implies we can conduct sampling under *q*(*G*) once, and reweight these samples for arbitrary values of *s* without having to conduct additional sampling. In other words, approximating *L*(*s*) using importance sampling does not require sampling under each value of *s* at which you want to approximate *L*(*s*).

In this paper we specifically consider the importance sampling proposal distribution 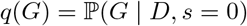, which corresponds to the posterior ARG assuming a neutral model. Later, we evaluate the performance of the estimator using the Markov chain Monte Carlo method ARGweaver, which samples from the posterior [37]. One can obtain the importance sampling estimate of the full likelihood *L*(*s*) by expressing Eq. 13 as an expectation over a different distribution, i.e. the posterior distribution of the ARG (assuming selective neutrality):

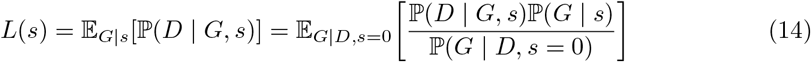

We can express Eq. 14 using the Monte Carlo approximation

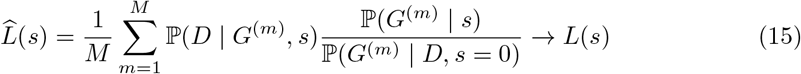

where 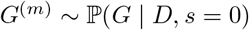, *m* = 1, 2,…, *M*, and “*→*”, here and in the following, means that the left-hand side converges almost surely to the right-hand side as *M* goes to infinity, assuming that a Law of Large Numbers for ergodic processes holds (the Birkhoff–Khinchin theorem).

Hence, if we sample genealogies from the posterior under selective neutrality, that is, 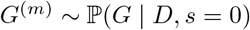, *m* = 1, 2,…, *M* (where *M* is the number of ARGs sampled), then the right-hand side of Eq. 15 can be used as a Monte Carlo estimator of the likelihood function. However, in practice this estimator is highly unstable. However, a more stable estimator of the likelihood ratio 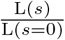 can be derived. We can divide through Eq.14 by 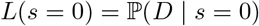 to get

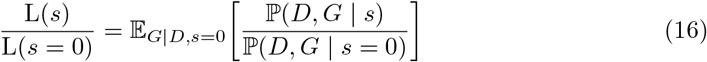

Because we assume the data are conditionally independent of selection given the full ARG, we can simplify this as

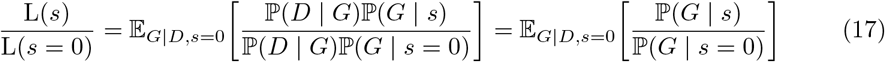

A key development in our method is that although we sample the ARG of the entire sequence, we only calculate likelihoods using the marginal tree at the selected site, which we will call *G*_*k*_. First, let us define *G*_*\k*_ as the rest of the ARG omitting the local tree at site *k*, *G*_*k*_. Consequently, *G* is equivalent to (*G*_*k*_, *G*_\*k*_).

We make a key assumption that, for differing sweep parameters *s* and *s′*

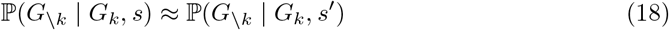

That is, we assume that *G*_\*k*_ is approximately conditionally independent of *s* given the marginal tree at the selected site, *G*_*k*_. Thus, we can reduce Eq. 17 to

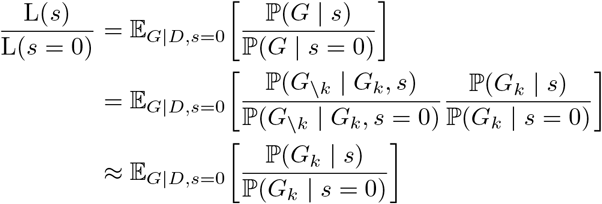

which suggests the following importance sampling estimator using genealogies sampled from ARGweaver will converge almost surely to a close approximation to the likelihood ratio:

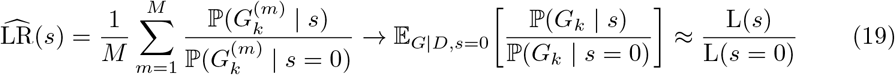

where 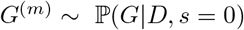 for *m* = 1, 2,…, *M*.

Finally, due to exchangeability of lineages within the derived and ancestral allelic classes, we can assume

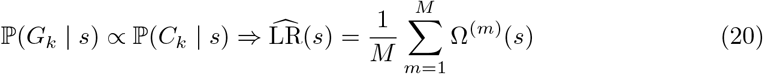

where

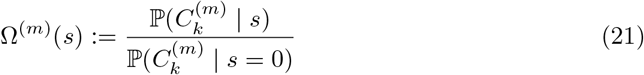

denotes the summand of the importance sampling estimator. That is, the topology within allelic classes is not important, and instead we need only the lines of descent process within each class.

We can maximize the likelihood ratio over different values of *s* to obtain the maximum-likelihood estimate of *s*

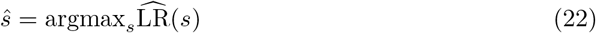

Finally, we show in S1 Text that an importance sampling estimate of *π*(*x*_*i*_ | *D, s*), the posterior marginal of the allele frequency at timepoint *i*, *X*_*i*_, is given by

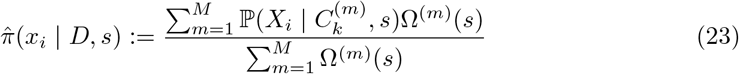

where in the summand we use the posterior marginal established in Eq. 11. In practice, we fix *s* = *ŝ*. A concern is, therefore, that this estimator does not take uncertainty in the estimate of *s* into account. This problem can be addressed by using a Bayesian approach, which we demonstrate briefly in S1 Text.

The method is implemented in package named CLUES, available for download at https://github.com/35ajstern/clues, with accompanying documentation currently provided in S1 Text. In this paper and the currect software release we assume positive directional selection with an additive effect on fitness, our method can be easily extended to general dominance relationships as well as negative selection.

### Simulations

To evaluate the power of `CLUES` to determine whether a site has been subject to selection, we simulated a dataset of *n* = 25 diploid individuals under two different demographic models; (1) a model of constant effective population size (*N* = 10^4^), and (2) a model of European (CEU) demography [43]. We performed both sets of simulations using the program `discoal` [44]. We set *µ* = 2*r* = 2.5 × 10^*−*8^ mut/bp/gen, *L* = 1 × 10^5^ bp or 2 × 10^5^ bp for the constant-size and CEU models, respectively, and simulated conditional on a variety of present-day frequencies and selection coefficients, the latter of which we ranged from weak to strong values. Under each condition, we simulated 100 independent iterations. We also sampled 1 ancient haplotype; because ARGweaver, which we used subsequently to sample the posterior ARG, does not incorporate any information about ancestral/derived states, it is best practice to add an ancient individual or outgroup to help polarize the the alleles. In practical settings where the ancestral state is unknown, ARGWeaver accomodates specification of missing data on the ancient haplotype. For the constant-size and CEU models, we used ancient sampling dates of 2 × 10^4^ and 1.6 × 10^4^ generations before present, respectively. Because `discoal` can only simulate piecewise-constant population sizes, we specified population sizes to take on the value of their harmonic mean over the epoch, calculated from the original CEU model. Commands to run simulations of trajectories, local trees, and haplotypes are described in S1 Text.

Importantly, we conditioned simulations on the site of interest segregating at a particular frequency in the present day. Hence, when we considered the power to discriminate between neutral and selected alleles, we controlled the present-day frequency to be equal in both of these cases. Avoiding this step would otherwise upwardly bias estimates of the statistical power, due simply to the tendency for selected alleles to segregate at higher frequencies than neutral alleles [45]. (If the allele frequency in itself is also of interest, this part of the likelihood could trivially be added at a later stage, by simply using the stationary distribution of the allele frequency; see “Allele frequency transition probabilities” in S1 Text.) We then simulate the allele frequency backwards in time, from the present-day frequency, until the allele reaches a frequency of 0. Simulators such as `discoal` achieve this by using the conditional Wright-Fisher diffusion (see e.g. [46]). In the case where effective population size changes over time, running conditional simulations requires additional considerations because the probability of a mutation entering the population scales approximately linearly with population size. Naively sampling the trajectory backwards in time will therefore produce a bias, unless trajectories where the mutation occurs while *N*_*e*_ is low are somehow penalized. Thus, approaches such as reweighting sample trajectories using importance sampling have been used to correct this bias [47]. The program `discoal` implements a similar bias-correcting scheme using rejection sampling that rejects trajectories where the mutation occurs while *N*_*e*_ is low with higher probability than trajectories where the mutation occurs while *N*_*e*_ is high.

Next, we inferred the posterior ARG given the sequence data we simulated using ARGweaver [37]. This method works by proposing adjustments to an initial ARG, and randomly accepting or rejecting these proposals based on calculations of the prior probability of the proposed ARG, as well as its likelihood given the sequence data. Because the prior probability is based on the effective population size, we specified the same effective population size in the prior as we used to generate the sequence data. We found it important to adjust the proposal mechanism of ARGweaver; specifically, we adjusted resample window size and the number of resamples per window to achieve an acceptance rate of about 30-70%. In total, we sampled 3 × 10^3^ ARGs for each simulation, discarding the first 1 × 10^3^ as a burn-in period, and subsequently thinning the remaining samples to reduce the computational burden of downstream analyses; we used a thinning rate of 100 samples, resulting in *M* = 20 approximately independent samples. Reducing the thinning rate would increase accuracy and convergence of the inference at the cost of additional computation to calculate the likelihood of each additional sample tree. Commands to conduct ARG-sampling and local tree extraction are described in S1 Text.

Using utilities in the ARGweaver package, we extracted local trees at the selected site (at the center of the locus) from these sample ARGs. We then analyze this final set of trees using `CLUES`. We also analyzed the same sequence data using nSL, *H*_12_, and Tajima’s *D* [14,19,48]. The nSL method is essentially equivalent to iHS [11], except nSL does not require specifying a genetic map; despite this, these methods have been shown to have very similar statistical power with a slight advantage of nSL under some conditions. *H*_12_ is a method to calculate haplotype homozygosity merging the two most common haplogroups; thus, it is a test for selection that is robust to the origin of a sweep, i.e. whether it is hard or soft. Tajima’s *D* is a site frequency spectrum-based statistic which is sensitive to skews in the frequency distribution of linked alleles caused by hitchhiking on the partially swept selected allele. We used scripts provided by [22] to calculate *D* and *H*_12_, using a window size of 100kb centered on the selected site. We compare testing for selection under these methods by comparing their power curves under both the constant *N*_*e*_ and CEU demography models (Figs. 3,4).

**Fig 3.**
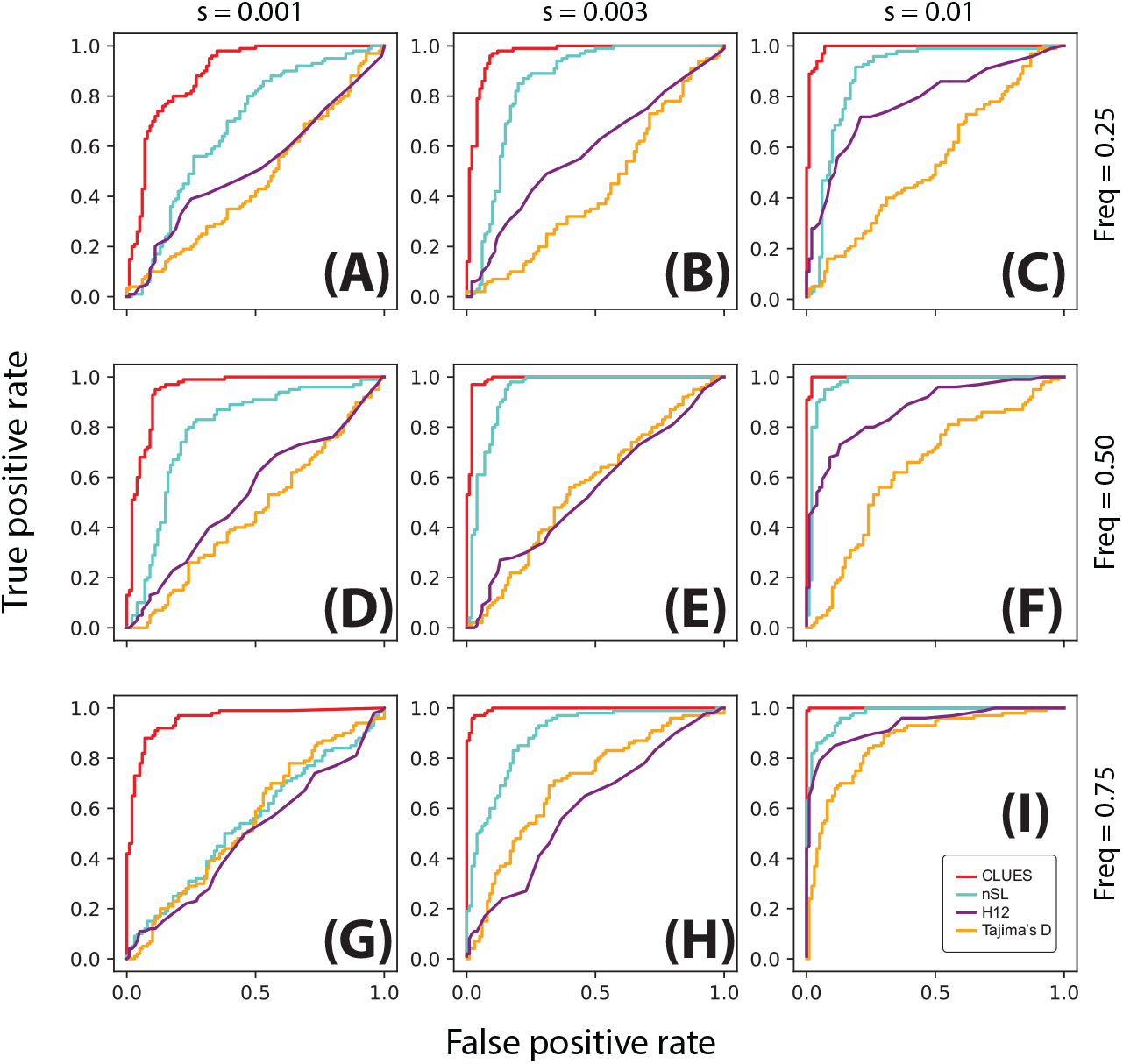
ROC curves illustrating performance of tests between selection and neutrality. Rows correspond to simulations conditioned on the same present-day allele frequency, and columns correspond to simulations with the same value of *s*. Simulations were performed under a model of constant effective population size (*N*_*e*_ = 10^4^) using a locus of 100kb, *n* = 25 diploid individuals and *µ* = 2.5 × 10^*−*8^ mut/bp/gen, *r* = 1.25 × 10^*−*8^ recombinations/bp/gen.

**Fig 4.**
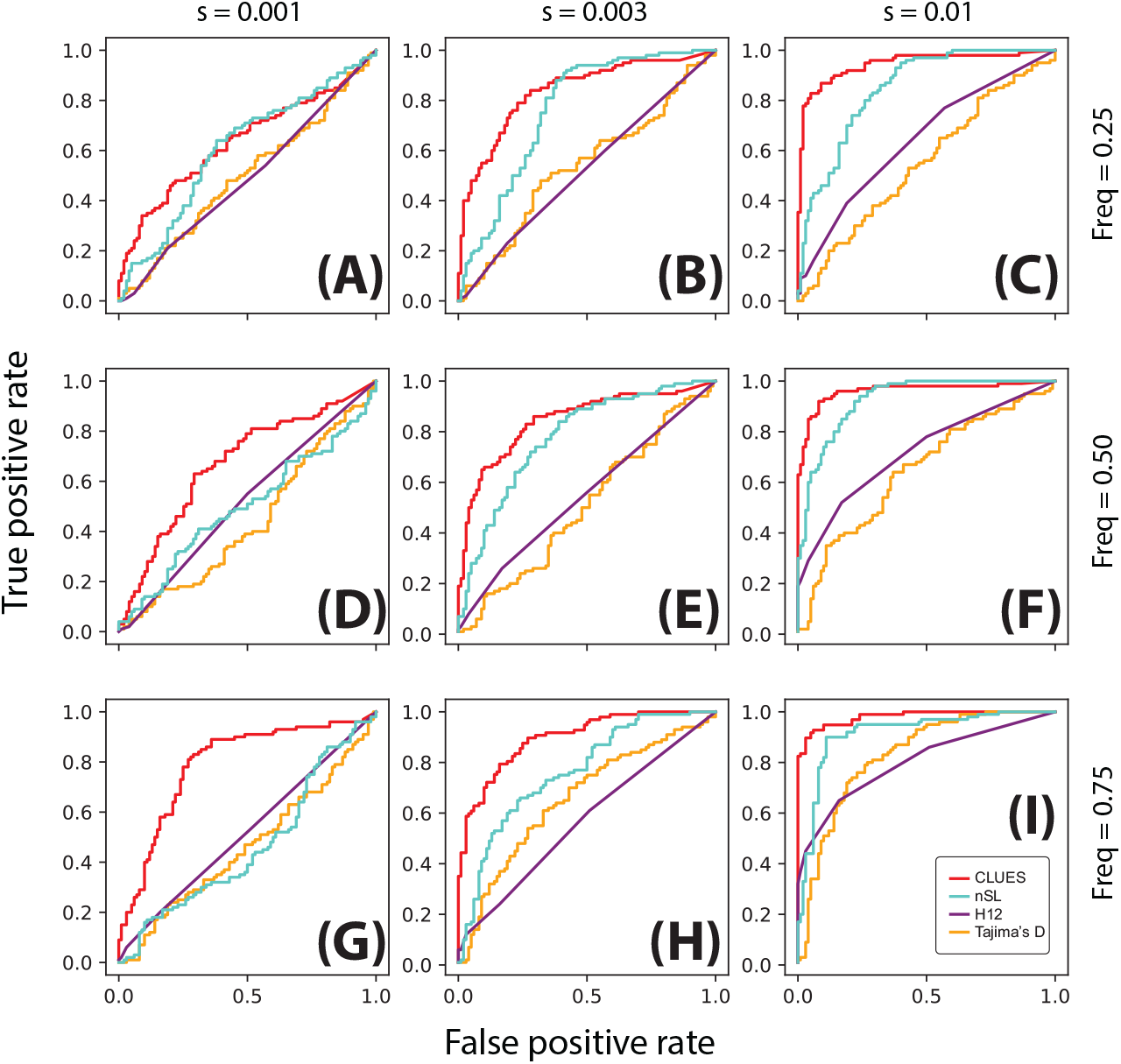
ROC curves illustrating performance of tests between selection and neutrality. Rows correspond to simulations conditioned on the same present-day allele frequency, and columns correspond to simulations with the same value of *s*. Simulations were performed under a model of European demography using a locus of 200kb, *n* = 25 diploid individuals and *µ* = 2.5 × 10^*−*8^ mut/bp/gen, *r* = 1.25 × 10^*−*8^ recombinations/bp/gen.

We also conducted a similar simulation study for detecting recent selection starting 100 generations ago. We simulated under the same CEU demographic model as previously described, but instead sample *n* = 50 diploids. We conducted ARG sampling and thinning as previously described, but in our analysis of the sample trees using `CLUES`, we calculated the likelihood for models of selection where *s* = 0 up until 100 generations ago, and *s* ≥ 0 from that point until the present day. This sweep from standing variation (SSV) model differs from the hard sweep model we used previously, which assumes *s* is constant throughout history. Instead of optimizing the likelihood function only with respect to *s*, we optimized with respect to two parameters, *s* and *t*_*s*_, jointly; here *t*_*s*_ represents the time of the onset of selection.

## Results

### Testing for selection

We found that across all scenarios, `CLUES` matches or exceeds the statistical power of the other methods evaluated (Figs. 3,4). As expected, all methods had highest power under large values of both the selection coefficient and the derived allele frequency (Fig 3I). Under these conditions, `CLUES` had 100% power at the 1% significance threshhold; the next most powerful method, nSL, had 68% power at the same significance level. `CLUES` also demonstrated improvement in power under weak selection; as the selection coefficient was decreased, nSL retained about 20% power when *s* = 0.003 and *<*5% power when *s* = 0.001, and Tajima’s *D* and *H*_12_ retained *<*5% power under both *s* = 0.001, 0.003 (Fig 3G,H). By contrast, `CLUES` retained approximately 45% and 90% power under *s* = 0.001, 0.003, respectively. We conclude that `CLUES` has high power across a wide regime of selection strengths, and has notably improved power over standard methods under weaker values of *s*.

We also considered the effect of present-day allele frequency on statistical power. Previous studies have shown a strong dependence of power on current allele frequency, with methods such as nSl and iHs having highest power at allele frequencies in the 70-90% range (see e.g. [11]). We tested for selection at alleles ranging in present day frequency from 25% to 75%, and while `CLUES` showed the expected pattern of increasing power with frequency, it also improved on the performance of other methods at lower frequencies. For example, under strong selection (*s* = 0.01), the power of `CLUES` changed from 100% to 90% to 85% as the frequency is decreased from 75% to 50% to 25% (Fig. 3C,F,I). By contrast, the power of the next most powerful method, *H*_12_, dropped from approximately 65% to 45% to 15% (Fig. 3C,F,I). Under moderate selection (*s* = 0.003), these effects were even more drastic, with the power of `CLUES` and nSL (the next most powerful method in this regime) changing from 90% to 60% to 50% and 20% to 5% to *<*5%, respectively. We conclude that `CLUES` has high power compared to standard methods across a wide range of allele frequencies, with the most major improvements in performance occurring when the derived allele is at lower frequencies (*<*50%). We found that using the approximation due to Griffiths (Eq. 6, [41]) decreased power of `CLUES` by increasing variability of the null distribution of the likelihood ratios. Hence, for testing under nonequilibrium demography we used the exact lines-of-descent probabilities (Eq. **??**). By contrast, as we will later show, we found the approximation given by Eq. 6 for *t ∈* [0, 1000] to improve estimation of allele frequency trajectories under this demographic model.

We also considered the same testing procedure under non-equilibrium demography, simulating under the previously described model of CEU demography (Fig. 34). We found in general reduced power to detect selection under this regime relative to the constant population size regime (Fig. 4I, cf. Fig. 3I), consistent with the well-known confounding of expanding population size with selection [10]. Nonetheless, `CLUES` demonstrated improved power relative to the competing methods across a wide range of selection coefficients (Fig. 4C,F,I), as well as across a wide range of derived allele frequencies (Fig. 4G,H,I).

### Estimating selection coefficients

Using the simulations from the previous section to study statistical power in testing for selection, we used our estimate of the likelihood surface for *s* to estimate the value of the selection coefficient via maximum likelihood (see Eq. 22). We obtained selection coefficient estimates under importance sampling using ARGweaver (Fig. 5), as well as selection coefficient estimates based on the true local tree observed directly (S1 Fig). Generally, the estimates are approximately unbiased. For example, the mean estimates of *s* = 0, 1 × 10^*−*3^, 3 × 10^*−*3^, 1 × 10^*−*2^ were approximately 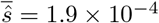, 9.6 × 10^*−*4^, 3.2 × 10^*−*3^, 1.3 × 10^*−*2^ when the present day frequency was fixed to 75% (Fig. 5A). Relative to inference when the true tree is observed, we found that the importance sampling estimates had increased variance, reflecting uncertainty in the tree. For example, we saw increased variability in the importance sampling vs. true tree estimates under constant population size (Fig. 5A vs. S1 FigA), as well as under CEU demography (Fig. 5B vs. S1 FigB). This pattern is consistent with the additional uncertainty in *s* when the local tree is not observed directly. Notably, we found that importance sampling under a model of CEU demography yields estimates with a slight bias towards lower values of *s*, especially under strong selection (e.g. *s* = 0.01).

**Fig 5.**
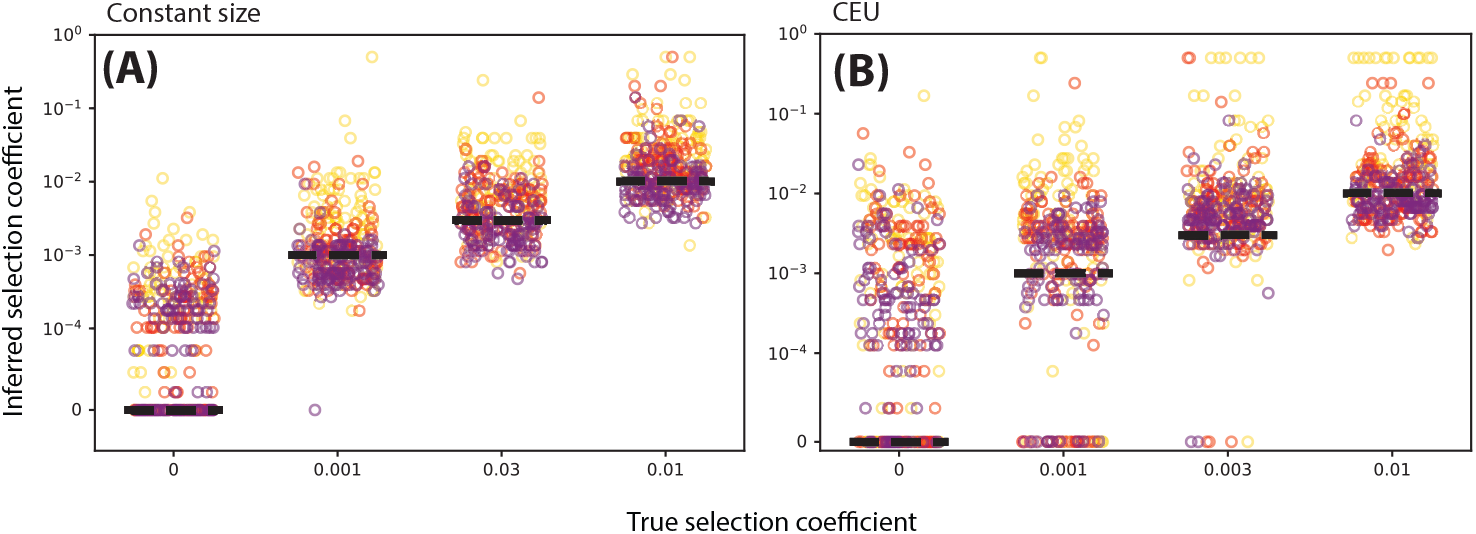
Inference of selection coefficients of varying strength using importance sampling method based on ARGweaver local trees. A: Constant population size. B: Tennessen CEU model. Marker color denotes present-day allele frequency (25/50/75% correspond to yellow/red/purple, respectively).

### Inferring allele frequency trajectories

Using the same simulations and importance sampling estimates we obtained in the previous sections, we decoded the hidden Markov model (HMM) described in the section Materials & Methods. Specifically, we take *ŝ*, the maximum likelihood estimate of *s*, and plug it into the posterior marginal (Eq. 11) to obtain a probabilistic estimate of the allele frequency during a particular epoch; we do this independently for each epoch in our discrete-time model. To get a point estimate, we choose to use the posterior marginal mean; i.e., for each epoch, we choose the mean of the posterior marginal distribution. We illustrate the accuracy of these allele frequency trajectory estimates assuming the true local tree is observed and under importance sampling when the true tree is unknown in Fig. 6. We find that estimates of the allele frequency trajectory are generally unbiased for both true trees (Fig. 6 A,B) and importance sampling (Fig. 6 C,D), with increased variance in the trajectory estimates in the importance sampling setting. We also illustrated variability in true vs. inferred trajectories controlling for *s* (S6 Fig, here setting *s* = 0).

**Fig 6.**
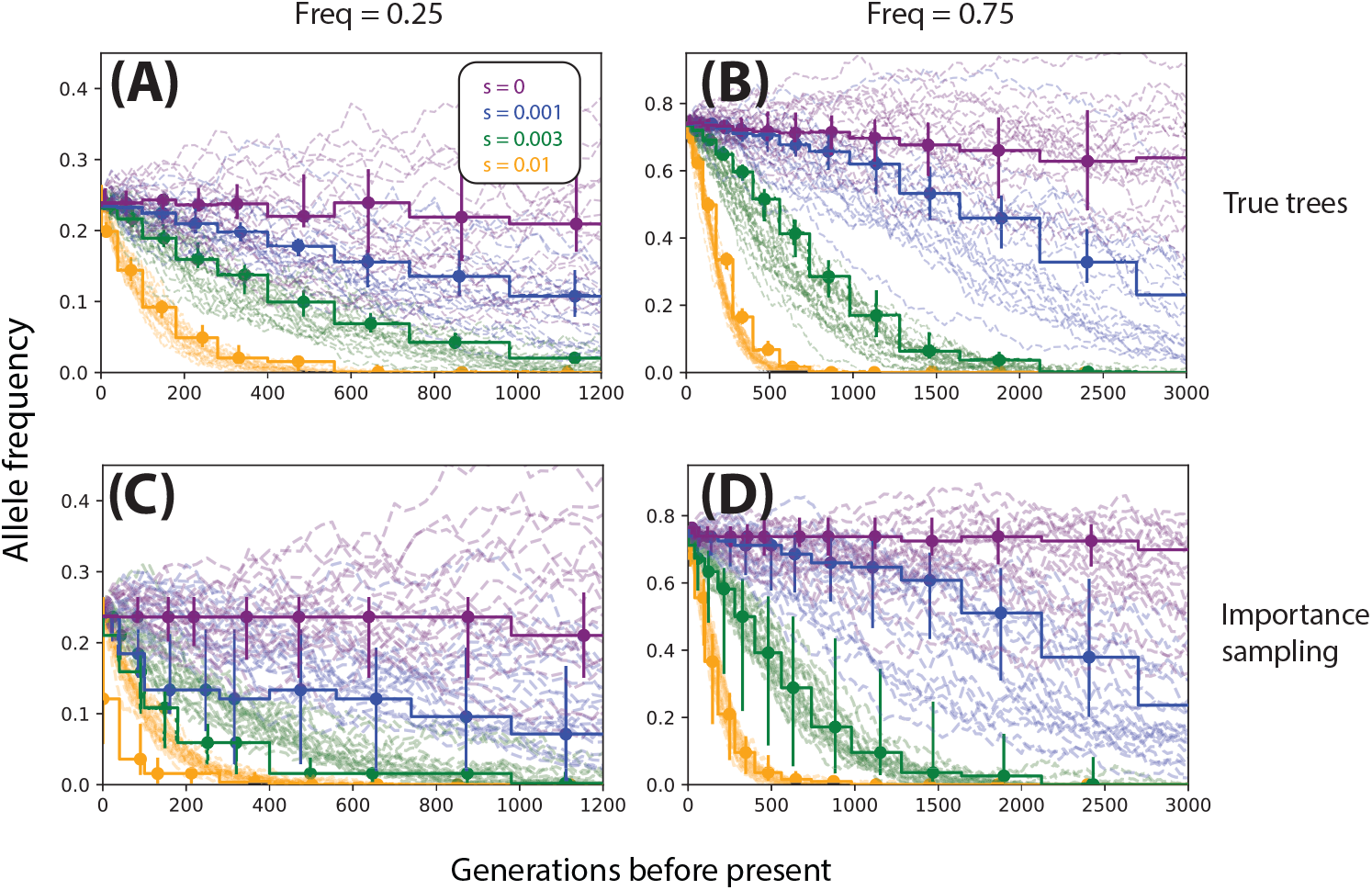
Allele frequency trajectories inferred from true trees (top row) and ARGweaver local trees (bottom row). Stepwise trajectories are inferred (vertical bars denote 25th-75th percentiles), dashed trajectories are the ground truth. Columns correspond to different initial allele frequencies (left: 25%, center: 50%, right: 75%), and rows/colors correspond to different selection coefficients. For each condition we show 25 randomly selected simulations and their corresponding inferences. All data are simulated under a model of constant effective population size (*N*_*e*_ = 10^4^).

Whereas inference tended to be relatively accurate for high-frequency alleles (Fig. 6 B,D), when the derived allele was simulated conditioned on lower frequencies (e.g. 25%, Fig. 6 A,C), estimates tend to be downwardly biased. We tracked this bias to a lack of convergence in ARGweaver; specifically, we found that across different demographic scenarios and selection coefficients, ARGweaver can drastically overestimate the occurrence of very recent coalescences (in our case, in the last 100 generations; see S5 Fig). Under constant population size, we see a nearly 7-fold excess in the number of recent coalescences inferred by ARGweaver. Naturally, this bias will affect estimates for low-frequency alleles more strongly, as fewer lineages subtend the derived allele, and thus a larger proportion of them are susceptible to this bias.

Because recombination rates vary substantially throughout the genomes of humans and other organisms, we also evaluated the accuracy of the estimates assuming *µ* = *r*, larger than the *µ* = 2*ρ* setting we used in the other simulations, and estimation accuracy to be robust to this increase in recombination rate (S2 Fig).

We also examined trajectory inference under non-equilibrium demography; i.e., the aforementioned model of CEU demography (S3 Fig). Under the CEU model, we found trajectory estimates to have increased variance under importance sampling vs. true trees, but also a slight downward bias in estimating the selection coefficient under strong selection (i.e. *s* = 0.01; see Fig. 5B, S3 Fig D). As this bias does not occur under the true trees (S1 Fig B, S3 Fig B), we inspected the posterior trees sampled by ARGweaver for patterns consistent with this bias. We found that under this demographic model in particular, ARGweaver tends to under-sample trees with short times to most recent common ancestor (TMRCAs; see S4 Fig). For reference, nearly 60% of runs under constant *N*_*e*_ contained even a single sample tree that had a TMRCA less than or equal to that of the true TMRCA (S4 Fig A). By comparison, under *s* = 0.01 and CEU demography, only 11% of ARGweaver runs met this criterion (S4 Fig B). Some bias is to be expected, as trees were sampled under a posterior distribution that assumes selective neutrality; however, these results suggest that, if ARGweaver is sampling from the true posterior assuming selective neutrality, then importance sampling estimates (of the selection coefficient, for example) will at least have much higher variance under the CEU model than under constant population size.

We further investigated whether uncertainty in *s* due to importance sampling variance drove the downward bias when estimating strong selection (Fig. 5B and S3 Fig D). First, we obtained importance sampling estimates of the trajectory fixing *s* to its true value (S7 Fig A). If uncertainty in *s* were the cause of the bias, then fixing the true value of *s* ought to correct for bias due to uncertainty. While we observe less bias in the estimates when fixing the true value of *s*, the bias is not totally eliminated. We observe a similar reduction in the bias of estimates under neutrality when we fix *s* = 0 (see S6 Fig B,E,H, vs. S6 Fig C,F,I). Thus, we conclude the bias is due to a lack of convergence in ARGweaver, which appears to be exacerbated in settings where strong selection is combined with non-equilibrium demography.

We also investigated whether incorporating uncertainty in the estimate of *s*, rather than fixing *s* = *ŝ*, would improve the accuracy of trajectory inference. One strategy for modeling uncertainty in *s* is to apply a prior distribution to *s*. We found that marginalizing out *s* with respect to its posterior distribution (assuming a uniform prior on *s*) did not have a noticeable effect on inference for large values of *s* (S7 Fig B). This result is concordant with our observation that for large values of *s*, the likelihood surface peaks so strongly that the posterior remains tightly concentrated around the MLE *ŝ*. Hence, applying a prior distribution to *s* does not appear to be an adequate strategy to model uncertainty in *s*.

### Inferring extremely recent selection

We applied our likelihood model of a sweep from a standing variant (SSV) to two types of datasets: selection from a standing variant starting 100 generations ago and selection with constant *s* (including *s* = 0), both described in ‘Simulations’ under Materials and Methods. We inferred trajectories under the best case scenario where the true trees are observed (Fig. 7A,B). We found that overall the method inferred the trajectory, as well as the strength and timing of selection, with highest accuracy when selection is strong (e.g. *s* = 0.03 in Fig. 7A,B). However, we found that as *s* took on smaller values (*s* = 0.01), many combinations of *s* and *t*_*s*_ had very similar likelihood (Fig. 7B), and thus estimates of *s*, *t*_*s*_, and the allele frequency trajectory tended to be noisier than under very strong selection (Figs. 7A,B). Adding the extra parameter *t*_*s*_ did not cause overfitting when inferring the trajectories of hard sweeps (Fig. 7A). We also found good power to distinguish between hard vs. soft sweeps (i.e. sweeps from a standing variant), as apparent in the trajectories inferred in Fig. 7A. We calculated statistical power to test for a hard sweep using the statistic 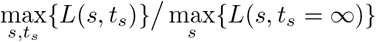; intuitively, this statistic is the ratio of the highest likelihood under any model with a SSV (*t*_*s*_ ≠ ∞) to the highest likelihood of any hard sweep (*t*_*s*_ = ∞). At the 1% significance level we found 60% and 100% power to distinguish soft vs. hard sweeps with *s* = 0.01, 0.03, respectively.

**Fig 7.**
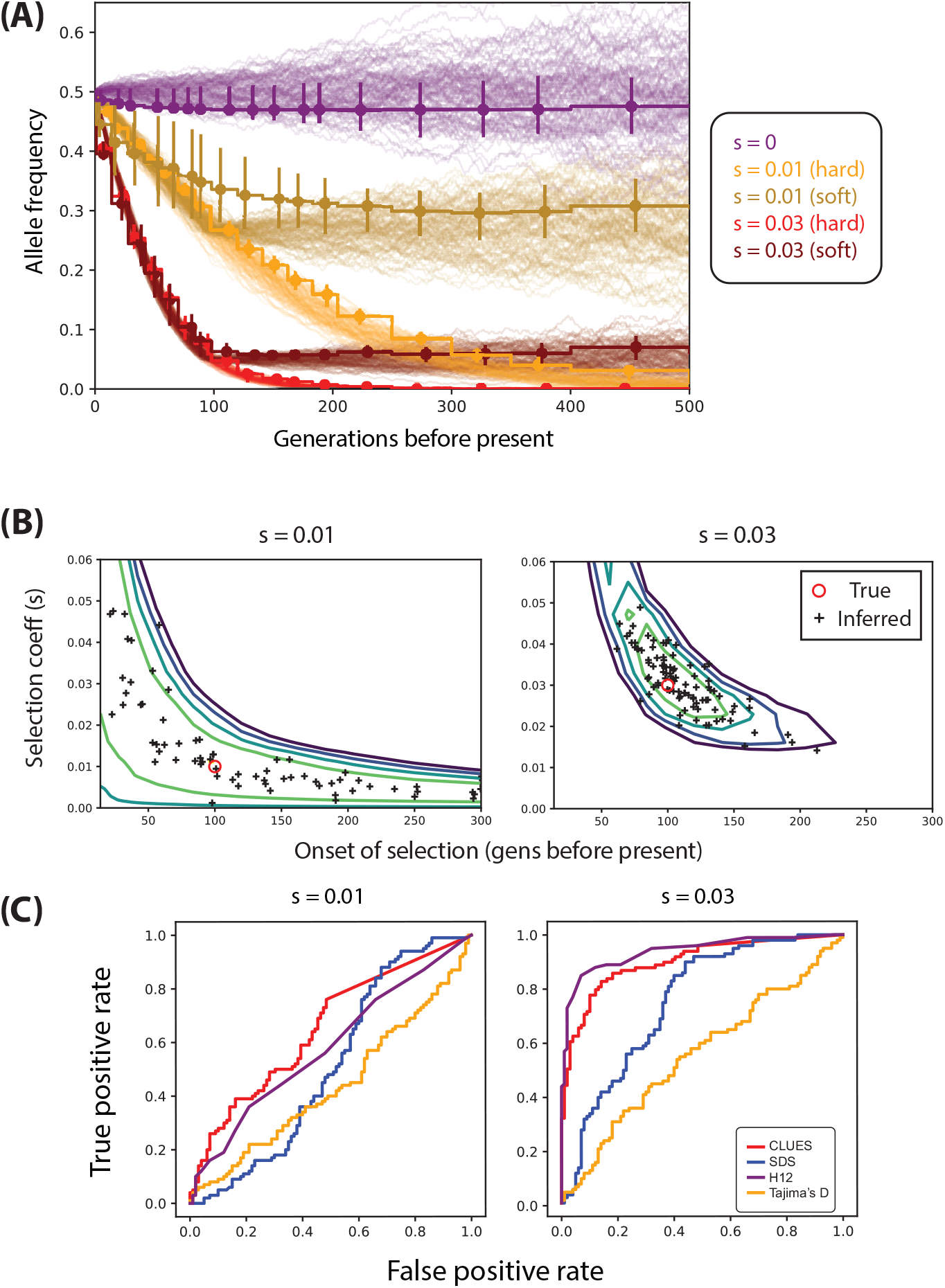
(A) Trajectories inferred from true trees under both hard sweeps and recent selection on a standing variant (i.e. soft sweeps) when both *s* and time of selection onset are unknown. (B) The log-likelihood surface for joint inference of *s* and onset of selection, averaged over 100 simulations, taking the true tree as observed. (C) ROC curves (using importance sampling) illustrating performance of tests between selection from a standing variant where onset of selection occurs 100 generations ago. We condition on a present day frequency of 50%.

We also performed importance sampling using ARGweaver and evaluated the power of the importance sampling estimates to detect recent selection vs. neutrality (Fig. 7C). Instead of comparing our method to nSL, which is not designed to detect signals of extremely recent selection, we compared to Singleton Density Score (SDS; [21]), as well as *H*_12_ and Tajima’s *D*. We found that for lower values of *s*, all methods had generally low power. Although `CLUES` exhibited fairly high power (44%) to detect very strong recent selection (*s* = 0.03) —even outperforming SDS—we found that *H*_12_ has about the same power (45%) in this particular case. The lower power (*<*5%) of SDS is consistent with the fact that the method was explicitly designed to have high power for large datasets (*n* > 1000 for selection coefficients of this magnitude). Although we demonstrate that `CLUES` has substantial power to detect extremely recent selection, we found that importance sampling point estimates of *s*, *t*_*s*_, and the trajectory were highly vulnerable to biases in the distribution sampled by ARGweaver (S5 Fig). Specifically, we found that across various demographic and selection conditions, ARGweaver samples trees with substantially more recent coalescent events than in the true trees. Specifically, under the European demographic model with the settings used here to study recent selection, we find ARGweaver samples about a 4-fold excess of recent coalescent events (S5 Fig B). Clearly, this bias would produce a false signature of recent selection under neutral conditions. Thus, we did not further explore importance sampling estimates of *s* and the trajectory under the recent selection model. We conclude that potential ARG-sampling methods that avoid this bias will improve upon power to detect recent selection, as well as point estimates of the strength, timing of selection, and the allele frequency trajectory.

### Fine-mapping the selected site

Strongly selected alleles tend to have high levels of LD to linked neutral sites, and thus many methods to detect selection are limited in their ability to determine the exact site under selection. To assess whether the likelihood ratio statistic produced by `CLUES` can be used to fine-map the selected site, we ran `CLUES` at linked neutral sites in a locus centered on a site under positive selection. Simulations were identical to those used in the simulation study under constant *N*_*e*_ = 10^4^, with a present-day selected allele frequency of 75%, *s* = 0.01, and 100 independent simulations. We chose sites with the maximal squared correlation coefficient *r*^2^ to the selected allele, such that *r*^2^ did not exceed a threshold value, and vary that threshold from 0.50 to 0.99 (Fig. 8). We found that when the true tree is observed (or sampled with high accuracy), the likelihood ratio statistic identifies the selected site correctly in a head-to-head test with the neutral linked site with 85% accuracy even when *r*^2^ ≤ 0.99; this quantity reaches 100% for *r*^2^ ≤ 0.50 (Fig. 8A). When the likelihood ratio is estimated using importance sampling via ARGweaver, the accuracy declines to about 50% and 85%, respectively (Fig. 8A). Because the exact causal site may not be known in many studies, we also investigated how the estimate of the selection coefficient, *s*, depends on *r*^2^ between the site analyzed and the site under selection. We estimate *s* given the true tree, and find that, on average, estimates of *s* decline with *r*^2^, such that for *r*^2^ ≤ 0.50, the mean estimate of *s* at these neutral sites is less than 20% of the true value of *s* at the causal site (Fig. 8B).

**Fig 8.**
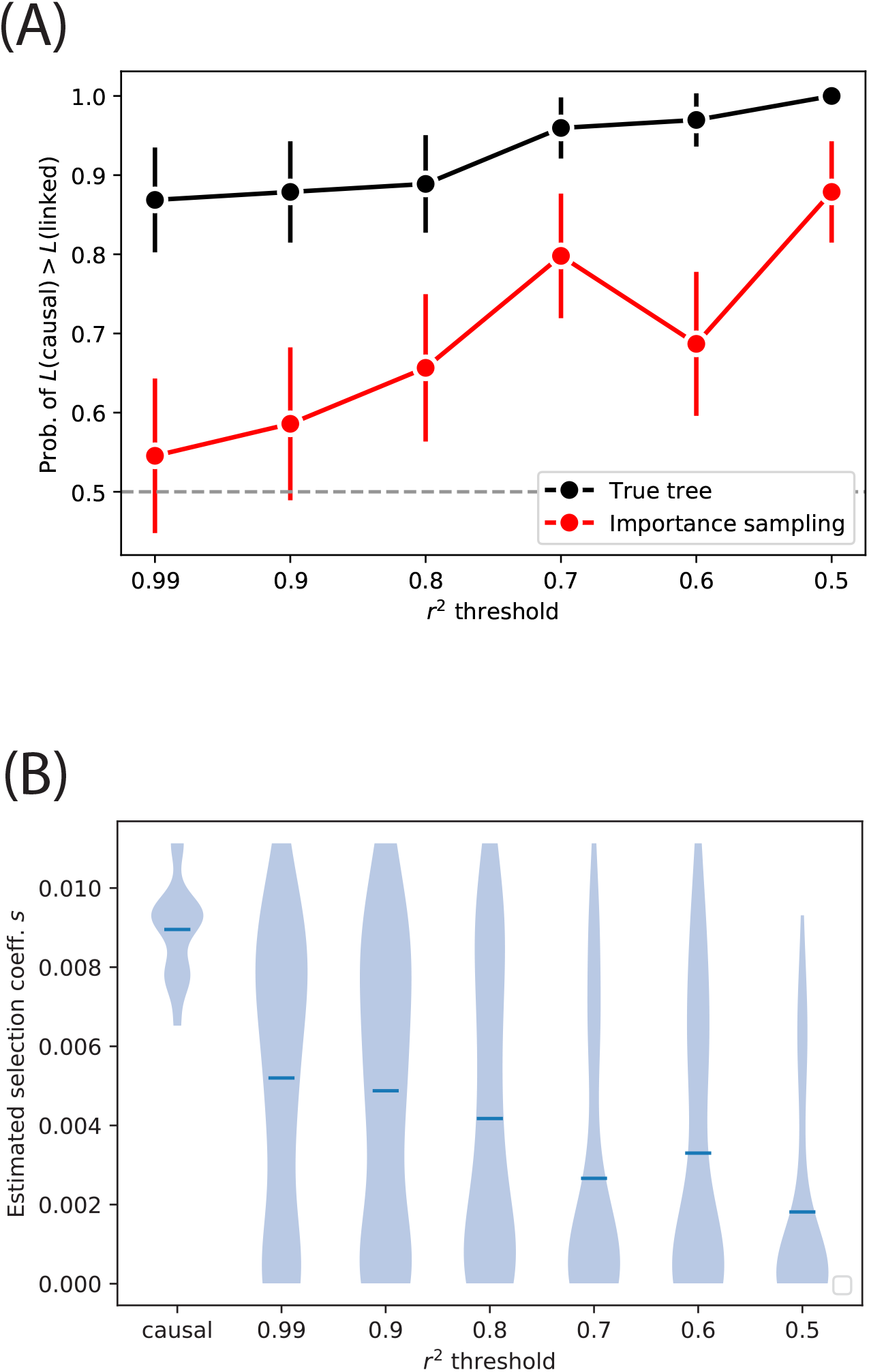
(A) Probability of correctly identifying the selected site in a head-to-head test with a linked neutral site. Vertical bars represent 95% CIs estimated by plug-in bootstrap. (B) Estimates of the selection coefficient at the causal vs. linked neutral sites, given the true tree. Mean estimates are represented by blue hash marks.

### Effects of background selection

We also assessed the effects of linked selection on inference of selection at a particular site. In particular, we consider the effects of background selection (BGS) on inference of positive selection on a linked site (S8 Fig). We performed forward simulations in SLiM 3 [49], assuming a model of constant *N*_*e*_ = 10^3^, a locus of length 1Mb, *n* = 25 diploid individuals and *µ* = 2.5 × 10^*−*8^ mut/bp/gen, *r* = 1.25 × 10^*−*8^ recombinations/bp/gen. We let mutations be neutral with probability 90% and (negatively) selected with probability 10%. We simulated under deleterious effects of *s* = 0*, −*0.001*, −*0.003 and −0.01, performing 100 independent replicates under each case. To simulate selection at a focal allele, 100 generations prior to the present day, we choose a random neutral allele conditional on its frequency falling in the interval [0.005,0.015] and its position falling in the interval [4 × 10^5^, 6 × 10^5^]bp (to ensure it is somewhat centered in the 1Mb region), and then endow it with a selection coefficient of *s* = 0.15. Prior to this timepoint, we perform a burn-in phase of 19900 generations with only neutral and deleterious mutation. From the sampled haplotypes, we perform importance sampling using ARGweaver to estimate the likelihood of the selection coefficient.

We find that our method is quite robust to BGS; e.g., the median estimate of the selection coefficient is approximately unbiased (mean *ŝ* = 0.13) as the strength of BGS is increased (S8 Fig). Also, regardless of BGS strength there is nearly perfect power to detect positive selection when comparing to neutral simulations (S8 Fig). We note that the strength of selection on the beneficial allele investigated here is somewhat strong; for weaker selection on a beneficial allele, inference with sites under BGS may be a more significant determinant of the beneficial allele’s trajectory (see e.g. [50]). We also note that our simulations assume a model fo equilibrium demography, but under non-equilbrium conditions (e.g., rapid population size expansion) BGS has a magnified effect on neutral diversity, which may further bias estimates of selection [51].

### Effects of demographic model misspecification

To explore the effects of demographic model misspecification on inference of selection, we ran `CLUES` on datasets simulated under a model of European demography (described earlier in Methods), using a mismatched model of constant *N*_*e*_ = 10^4^ to calculate the likelihood, and compare them to calculations under the true demographic model (S9 Fig). We report likelihood ratios (S9 Fig A,C) as well as estimates of the selection coefficient *s* (S9 Fig B,D) both given the true tree (S9 Fig A,B) and approximated via importance sampling using ARGweaver (S9 Fig C,D). We find that statistical power to detect selection, especially strong selection, is not substantially impeded by model misspecification (S9 Fig A,C). We do, however, find upward bias in estimates of the selection coefficient when *s* is 0 or close to 0 (S9 Fig B,D).

### Analysis of a lactase persistence SNP

To assess performance of `CLUES` on empirical data, we applied our method to study selection acting on the SNP rs4988235 in the *MCM6 gene*, known to regulate the neighboring *LCT* gene and affect the lactase persistence trait. The derived allele (A) current segregates at approximately 72% in the 1000 Genomes Phase 3 reference panel (British in England and Scotland, henceforth GBR; see S10 Fig A). We conducted sampling in ARGweaver assuming a model of European demography [43], using a 300kbp region centered around the focal SNP and polarizing alleles using the genomes of three ancient individuals (Altai Neandertal, Denisova, and Vindija Neandertal [52–54]). We sampled *M* = 200 ARGs, extracted local trees using tools in the ARGweaver package, and conducted importance sampling to estimate likelihood surfaces and trajectories using `CLUES`.

We found very strong evidence for selection on rs4988235 (*s* = 0.0161, log LR = 131.82). The trajectory as well as the value of the selection coefficient inferred by `CLUES` are consistent with previous estimates of the trajectory and *s* = 0.018 due to Mathieson and Mathieson (2018), illustrated in Fig. 9 [55]. Their method incorporates genomic times series spanning thousands of generations using an HMM-based approach, where hidden states are population-wide allele frequencies, observed states are genotypes of sampled ancient individuals, and transition probabilities are governed by the selection coefficient. Our approach, by contrast, does not utilize any ancient/timecourse data except for the 3 aforementioned ancient individuals, which we use to simply polarize the derived and ancestral states of each allele.

**Fig 9.**
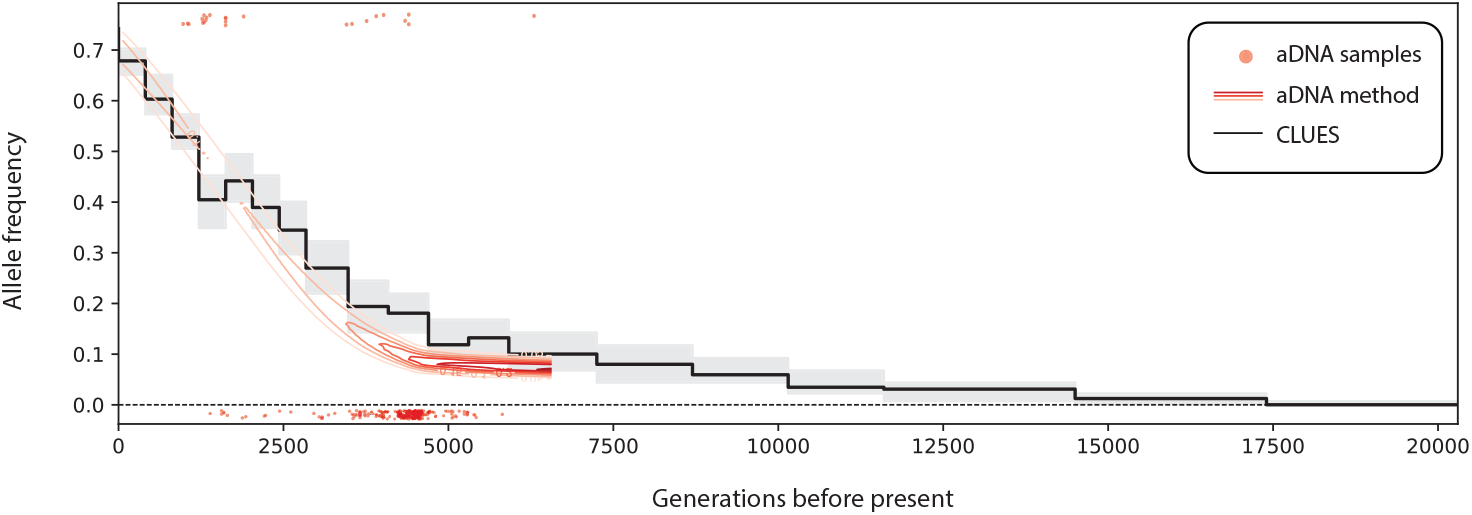
Comparison of inferred allele frequency trajectories for a sweep at rs4988235 (MCM6) in GBR under an ancient DNA (aDNA) based method vs. `CLUES`, which only uses contemporary modern data. Black curve is the posterior median allele frequency, whereas gray areas are a 95% posterior interval. The red surface is the posterior of the frequency trajectory within Steppe ancestry conditioned on an ancient DNA time series, adapted from [55].

### Analysis of *EDAR*

Another canonical example of a common SNP under selection in humans in rs3827760 (see e.g. [56]). This SNP codes for a V*→*A change that is present at high frequency in East Asian populations (e.g. 94% in Han Chinese [CHB] in the 1000 Genomes database), intermediate-high frequency in Central and South America, and low frequency in other geographical regions S10 Fig. This variant is associated with a number of traits, including tooth shape and hair straightness [57,58]. To estimate selection on this SNP, we conducted sampling in ARGweaver assuming a model of East Asian demography [59], using a 300kbp region centered around the focal SNP and polarizing alleles using the genomes of three ancient individuals (Altai Neandertal, Denisova, and Vindija Neandertal [52–54]). We sampled *M* = 200 ARGs, extracted local trees using tools in the ARGweaver package, and conducted importance sampling to estimate likelihood surfaces and trajectories using `CLUES`. We estimate that rs3827760 has undergone selection with *s* = 0.0047, corresponding to an allele age of roughly 45kya (S11 Fig). Our estimates are in stark contrast to some previous estimates obtained using ABC methods, which estimate *>* 30-fold stronger selection on this SNP, and an allele age of 1.4-6.9kya [32]. Our results are consistent with ancient DNA evidence, which suggest the derived allele to have originated prior to 7.5kya [8].

### Analysis of pigmentation alleles

Using the same GBR panel from 1000 Genomes Phase 3, we analyzed a set of SNPs associated with pigmentation-related traits, some of which were previously identified as likely targets of recent selection [21]. We conducted sampling in ARGweaver assuming a model of European demography, using a 300kbp region centered around the focal SNP and sampling *M* = 200 approximately iid ARGs. We ran `CLUES` and estimated likelihood surfaces and allele frequency trajectories for these SNPs (Fig. 10). We found significant concordance between the SDS values and our likelihood ratio statistics paired for each SNP (*p* = 1.7 × 10^*−*3^, Spearman one-sided) [21]. We also illustrated the geographical distribution of these SNPs among diverse populations (S12 Fig) using GGV [60].

**Fig 10.**
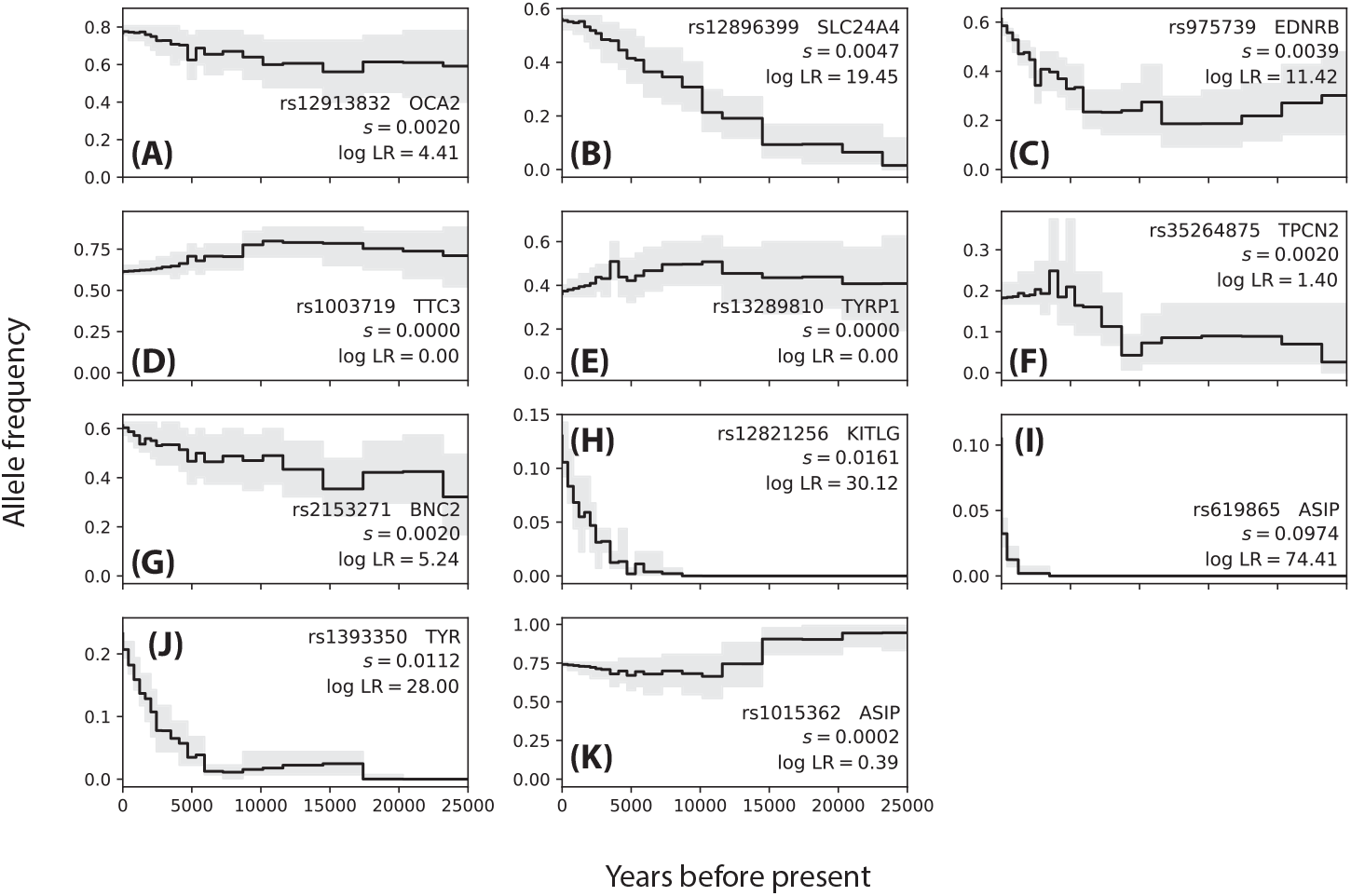
Allele frequencies trajectories inferred for 11 pigmentation-associated SNPs in GBR.

We found several signals of very strong selection acting on rs619865 (*ASIP*, *s ≈* 0.10, Fig. 10I), rs12821256 (*KITLG*, *s ≈* 0.016, Fig. 10H), and rs1393350 (*TYR*, *s ≈* 0.011, Fig. 10J); these SNPs are significantly associated with freckling, blonde hair color, and freckling and blue/green eye color, respectively [61–63]. Interestingly, these SNPs all demonstrated a signal of selection mostly concentrated in the last *∼*5 kya. The geographical distribution of the frequency of these SNPs shows that the derived version of these variants are mostly concentrated in European populations, with minimal sharing with populations located in Africa and Asia (S12 Fig I,H,J). For example, *TYR* and *KITLG* segregate at a frequency *∼*20% in several European populations and have a frequency close to 0% in African and East Asian populations (S12 Fig J). These three SNPs are the only ones in this set of SNPs which have a frequency of nearly 0% across the African populations surveyed, with the exception of OCA2/HERC2 (S12 Fig A,H,I,J), consistent with our evidence for recent selection at these loci. The frequencies of these variants in GBR ranges from *∼*10-20%; by contrast, the only other variant in this set with comparable frequency in GBR (13%), rs35264875 (*TPCN2*), we find inconclusive evidence of selection (Fig. 10F), consistent with its comparably even geographical distribution relative to the aforementioned SNPs at *ASIP, KITLG*, and *TYR* (S12 Fig F).

At rs12896399 (*SLC24A4*, Fig. 10B), a SNP identified to be significantly associated with hair color [62], we found strong evidence for moderate selection (*s* ≈ 0.005). This result is consistent with a previous analysis that suggested positive selection acted on this allele in Out-of-Africa (OoA) populations, based on its high allele differentiation relative to a YRI panel, and low haplotype diversity within CEU individuals [63]. Our results, paired with the apparent low levels of differentiation between European and Asian populations relative to differentiation between OoA populations and African populations at this locus (S12 Fig B) are consistent with our estimate that selection acted on *SLC24A4* as early as *∼*30 kya, during the OoA bottleneck as inferred by [43,59].

Notably, we find moderate evidence for selection on rs12913832 (*OCA2/HERC2*, Fig. 10 A, S13 Fig), a SNP previously shown to be causal for blue-brown eye color [64] and significantly associated with hair color [62]. This gene exhibits abberantly high differentiation across populations [65], consistent with a model of local adaptation of eye color. Compared to previous estimates based on ancient DNA samples [66], we estimate substantially weaker selection acting on this gene (*s* ≈ 0.002 vs. *s* ≈ 0.04), and we find no evidence to support a recent increase in selection acting on this SNP (i.e., our method found a hard sweep to have higher likelihood than a SSV). Our estimate of moderate selection and lack of a recent change in the selection coefficient imply that selection on OCA2/HERC2 began at least *∼*50 kya, roughly the time of the start of the OoA bottleneck estimated by [43,59]. Our analysis suggests that selection on *OCA2/HERC2* may have begun much earlier than previously suggested [66]. We also note that the aforementioned rs12913832 (*OCA2/HERC2*)—as well as rs2153271 (*BNC2*), a SNP which is significantly associated with freckling (Fig. 10 G)—occur in high-frequency archaic haplotypes [67]. While our method is not explicitly designed to control for population structure between archaic and modern human lineages, we do find moderate evidence for selection on both of these SNPs.

One surprising result is that we found no signal of selection acting at rs13289810 (*TYRP1*, *s* ≈ 0, Fig. 10E). In Europeans, *TYRP1* is associated with hair and eye pigmentation [68–71]. Some analyses of European populations have indicated evidence for positive selection on *TYRP1* [56,63,69]. Our results temper these claims, and appear consistent with the fairly even geographical distribution of rs13289810 frequency across European, African, and Middle Eastern populations (S12 Fig E).

## Discussion

We have developed an approach to use modern population genomic data to approximate the full likelihood of selection acting on a locus. We use this approach to test for and estimate the strength and timing of selection, as well as estimate the full allele frequency trajectory. The method is effective across a span of selection coefficients (*s* = 0 *−* 0.01), derived allele frequencies (*f* = 25% *−* 75%), and under multiple demographic models.

Our method draws on previously published methods to estimate the ancestral recombination graph (ARG). We chose to use ARGweaver because it is the only currently available method for sampling the posterior of the ARG; as shown in our derivation of the importance sampling estimates, we rely on sampling from the posterior in order to make rigorous guarantees regarding convergence and consistency of our estimators. Intuitively, it is important to model the uncertainty in the local tree in order to marginalize out this latent variable. We showed that estimates of the selection coefficient and the trajectory are generally accurate, barring scenarios where importance sampling is inefficient, or ARGweaver produces a bias in the inferred trees. In light of these biases, under certain conditions—primarily when the derived allele is at low frequencies (≤ 25%)—importance sampling using ARGweaver trees has limited power to detect selection.

Another important limitation of ARGweaver is its computational cost; in order to study selection on short timescales, large sample sizes are necessary, often on the order of thousands of individuals [21]. The runtime of ARGweaver grows dramatically with increasing sample size; not only does the cost of the individual sampling steps increase with sample size, but also so does the size of the state space, necessitating more samples be taken in order to achieve convergence to the stationary distribution.

However, we see potential to make use of recent advances in inference of local trees in order to further advance approximate full-likelihood methods to infer selection (see e.g., [72–75]; it is worth noting that some of these methods, such as [75], do not infer the ARG in a strict sense, but rather the sequence of local trees along a recombining locus). A major benefit of these methods is that they are far more scalable than ARGweaver, and hence offer more potential to study selection on short, punctuated timescales. However, they also possess several limitations: Firstly, several of these methods only infer topologies, rather than branch lengths [73,74]. While it is possible to infer branch lengths condition on topology estimates, it is unclear how accurate these estimates would be. By contrast, methods that infer branch lengths along with topology entail a slight tradeoff in their scalability [72,75]. Another limitation of these methods it that they only yield a point estimate of the local tree, rather than estimating uncertainty in the tree. Nonetheless, it may be feasible to quantify uncertainty in the local tree using a jackknife approach where the local tree is inferred over random subsets of the individuals.

It may also be possible to make use of recent advances in inferring pairwise coalescence times (e.g., [76]) to build an approximation to the full likelihood. Recently, Albers & McVean proposed a composite likelihood method to estimate allele age by “sandwiching” the age using identity-by-descent tracts at the site of interest [77]. However, their method does not extend to inferring how the allele frequency changed over time, and does not explicitly model selection.

Currently our method assumes correct knowledge of the demographic history. The effects of latent or mis-specified population structure on inference of selection are well known (e.g., [78]), but in future work one might try to determine the exact effects of mis-specification of effective population size on both inferring the local tree, and inferring selection conditional on the local tree. One approach to dealing with this is to extend the importance sampling approach we use to correct for selection to additionally correct for demography, when ARG sampling is performed under a mis-specified demographic model.

Furthermore, many aspects of our model of selection (e.g. coalescence, allele frequency transitions) assume a panmictic population. To extend our model to more complex demographic models and/or linked selection (i.e., allowing multiple sites to be subject to selection) would entail drastically increased computational cost (e.g., marginalizing allele frequencies corresponding to each population, rather than the allele frequency in a single population). Using a deterministic approximation of the allele frequency trajectory would circumvent this issue, but it would also raise new issues, such as how to model allele frequencies when *s* = 0.

Despite its limitations, the method presented here provides the first close approximation to a full likelihood function for the selection coefficient under simple models. As demonstrated by our simulations, full likelihood methods have the potential to greatly improve power to detect selection and estimate the strength of selection under a variety of conditions. It also provides a rigorous and accurate method for estimating allele frequency trajectories, and is the first to achieve so using modern data. As methods for inferring ARGs improve in the future, so too will the derived methods for detecting and quantifying selection and inferring allele frequency changes.

## Acknowledgments

We thank Melissa Hubisz and Andrew Kern for extensive help with the software packages ARGweaver and `Discoal`, respectively. We also thank Graham Coop, Michael Edge, Yun Song, as well as Vladimir Shchur and other members of the Nielsen Lab, for helpful discussions.

## Author contributions

AJS: Conceptualization, formal analysis, investigation, methodology, software, validation, visualization, writing (original draft), writing (review and editing); PRW: Software, writing (original draft); RN: Conceptualization, methodology, supervision, writing (original draft), writing (review and editing).

## Supporting information

**S1 Fig.**
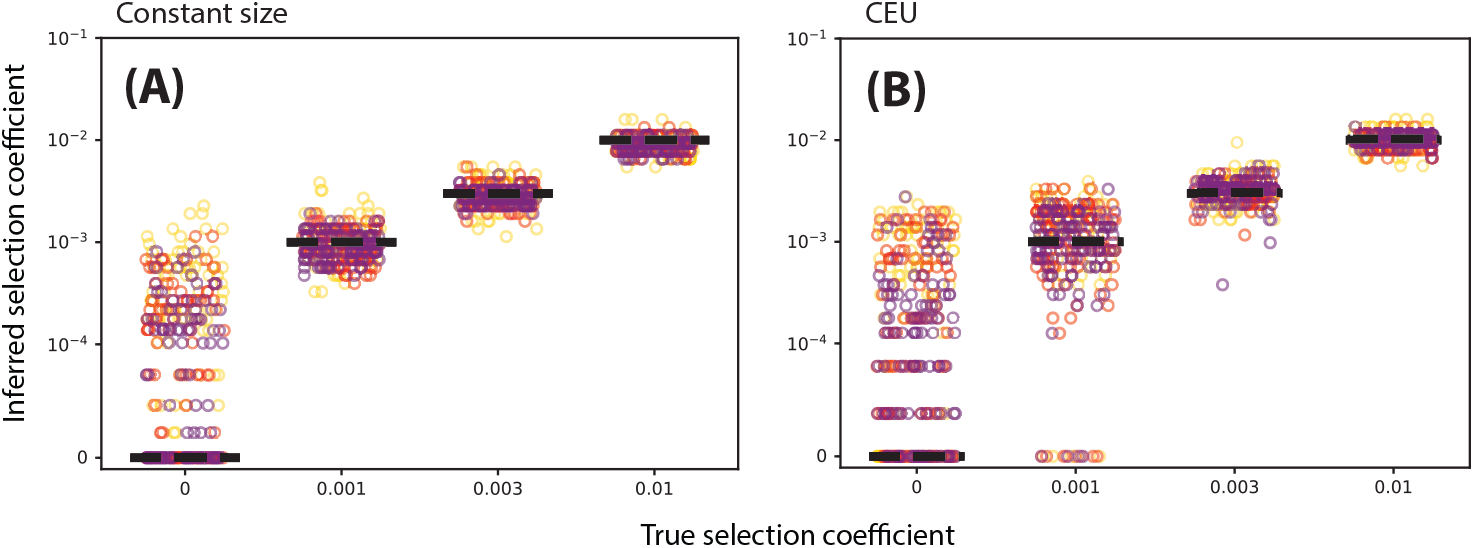
Selection coefficients inferred directly from the true local trees. Left: constant population size (*N*_*e*_ = 10^4^). Right: Tennessen CEU demographic model. Marker color denotes present-day allele frequency (25/50/75% correspond to yellow/red/purple, respectively).

**S2 Fig.**
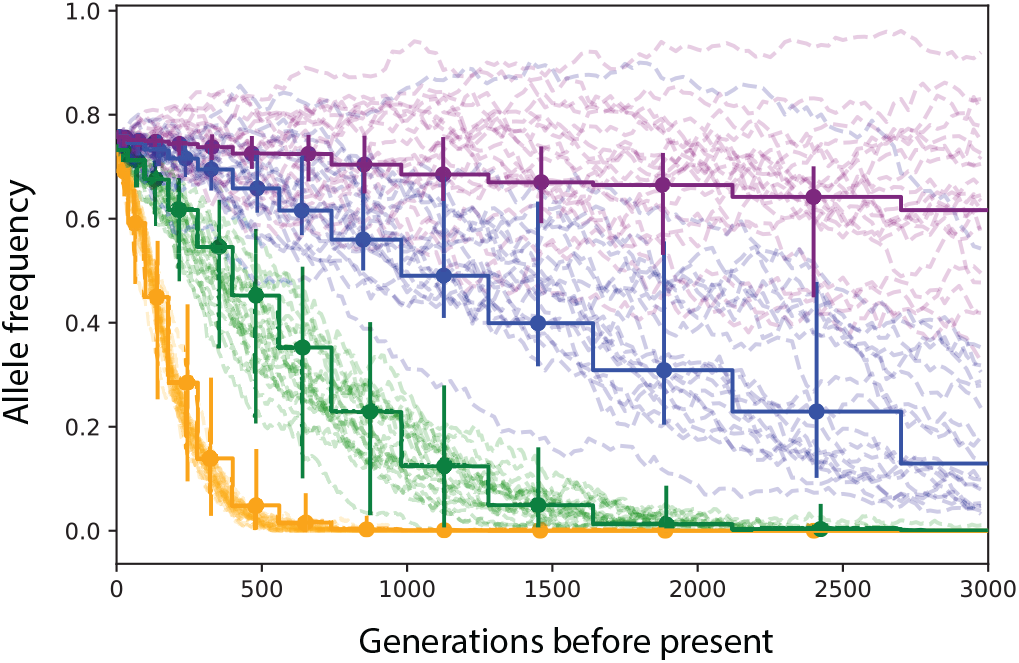
Allele frequency trajectories inferred from ARGweaver local trees when *µ* = *r*. We set *µ* = 2.5 × 10^*−*8^ mut/bp/gen, *r* = 2.5 × 10^*−*8^ recombinations/bp/gen and fix the present day allele frequency to *X*_0_ = 50% Stepwise trajectories are inferred, dashed trajectories are the ground truth. Vertical bars denote the 25-75th percentile range of estimates. For each condition we show 20 randomly selected simulations and their corresponding inferences. All data simulated under a demographic model with constant size *N* = 10^4^.

**S3 Fig.**
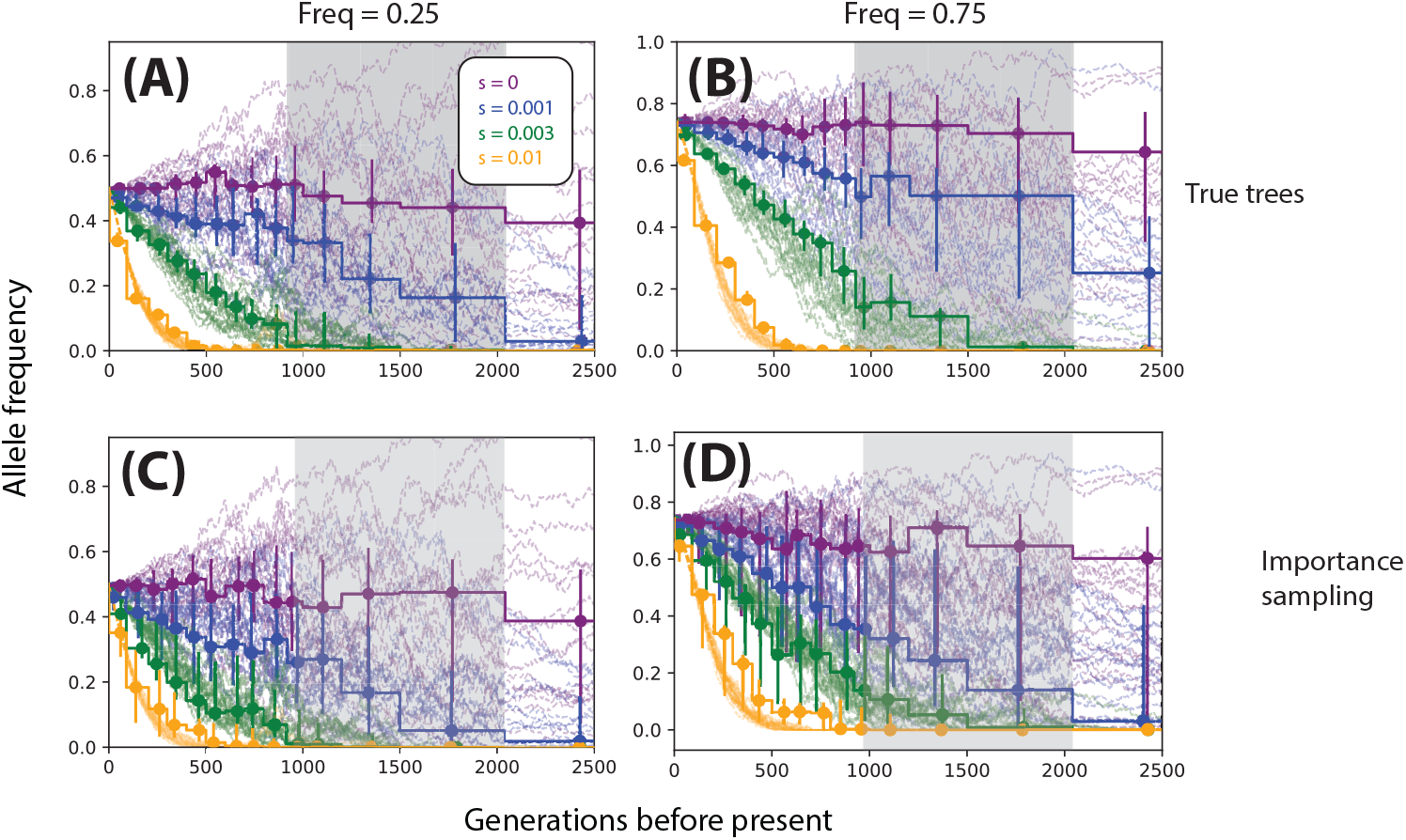
Inferring allele frequency trajectories under CEU demography. Trajectories were inferred from true local trees (top row) and importance sampling on ARGweaver local trees (bottom row). Columns correspond to different present-day allele frequencies (left: 50%, right: 75%). For each condition we show 20 randomly selected simulations (dashed, translucent lines) and their corresponding inferences (piecewise constant curves; dots and vertical bars indicate the median and 25-75 percentiles of estimates, respectively). The gray box indicates the timing of the bottleneck, occurring approximately 920-2040 generations ago. Simulations were done under the European demographic model described in Methods and Materials using a locus of 200kb, *n* = 25 diploid individuals and *µ* = 2.5 × 10^*−*8^ mut/bp/gen, *r* = 1.25 × 10^*−*8^ recombinations/bp/gen.

**S4 Fig.**
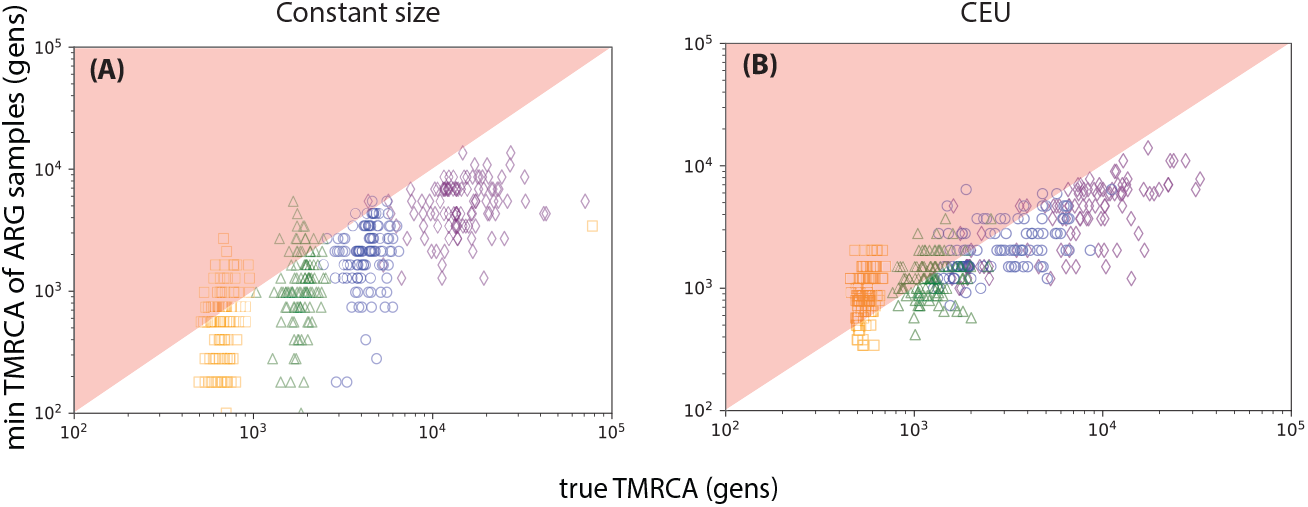
ARGweaver proposes less accurate trees under non-equilibrium demography. Left: constant *N*_*e*_ = 10^4^. Right: Tennessen CEU demographic model. We found in Fig. S2 Fig that importance sampling using ARGweaver tends to underestimate the selection coefficient under a model of CEU demography. To demonstrate that the proposal distribution for sampling the local tree is the source of this bias, we use TMRCA of the local tree as a heuristic the locus’s selection coefficient. For the sake of argument, we postulate that as one decreases the TMRCA of a local tree, the maximum-likelihood estimate of the selection coefficient strictly descreases. If so, then if the minimum value of the sampled TMRCAs is greater than the true TMRCA, then this instance of the importance sampling estimate will underestimate the selection coefficient. Hence, one can measure importance sampling efficiency by looking at the probability that the minimum value of the sampled TMRCAs is less than the true TMRCA. This is shown graphically by the proportion of points that fall in the red upper triangle. The selection coefficients *s* = 0, 0.001, 0.003, 0.01 are indicated by purple, blue, green, and orange, respectively. Simulations were done under the European demographic model described in Methods and Materials using a locus of 200kb, *n* = 25 diploid individuals and *µ* = 2.5 × 10^*−*8^ mut/bp/gen, *r* = 1.25 × 10^*−*8^ recombinations/bp/gen. We fixed the present-day allele frequency to be 75%.

**S5 Fig.**
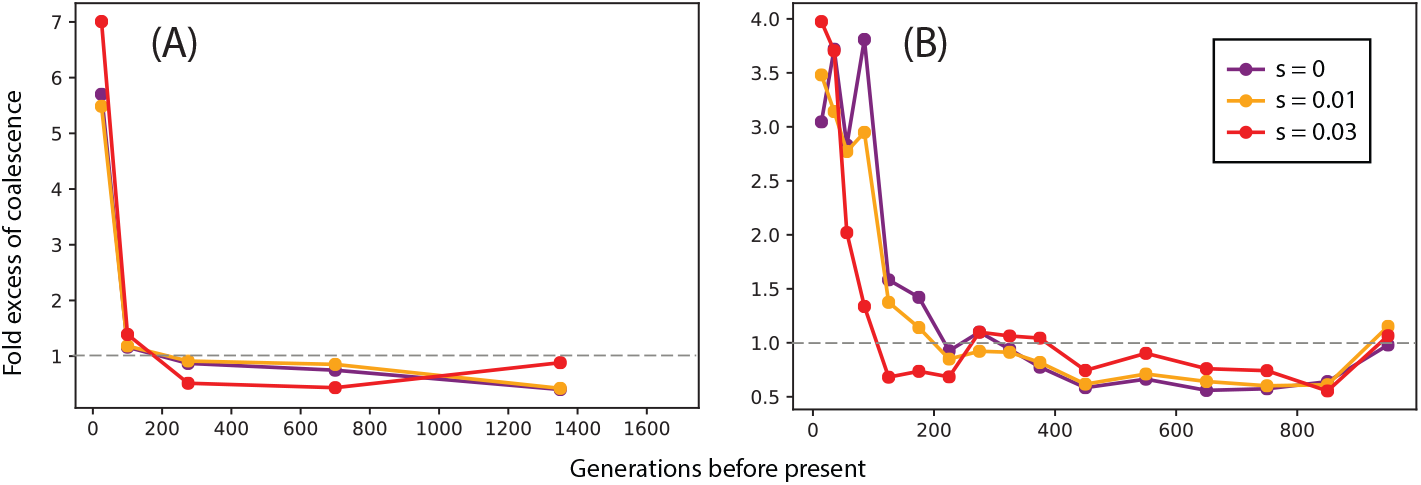
ARGweaver infers an excess of recent coalescences. As a diagnostic for the trees outputted by ARGweaver, we compared the amount of coalescence in the sample trees vs. the local trees. Let *a*_*i,s*_ be the number of coalescent events during epoch *i* in the *s*th replicate of the simulation. We calculate

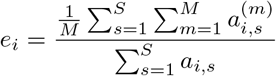

as an estimate of the “fold excess” of coalescence, where 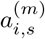 denotes the *m*th ARGweaver sample of *a*_*i,s*_. Notice that if the sample trees closely approximate the true tree, then *e*_*i*_ ≈ 0. The dashed line indicates no excess, i.e., no bias in the estimates. We find that ARGweaver can have a nearly 4× excess of inferred coalescence events in the most recent epochs (e.g. [0,100] generations ago). Here we simulated recent selection starting 100 generations ago under a model of European demography with *n* = 50 diploid individuals, a physical length of 200kb, and *µ* = 2.5 × 10^*−*8^ mut/bp/gen, *r* = 1.25 × 10^*−*8^ mut/bp/gen. We condition on the variant having a present-day frequency of 50%.

**S6 Fig.**
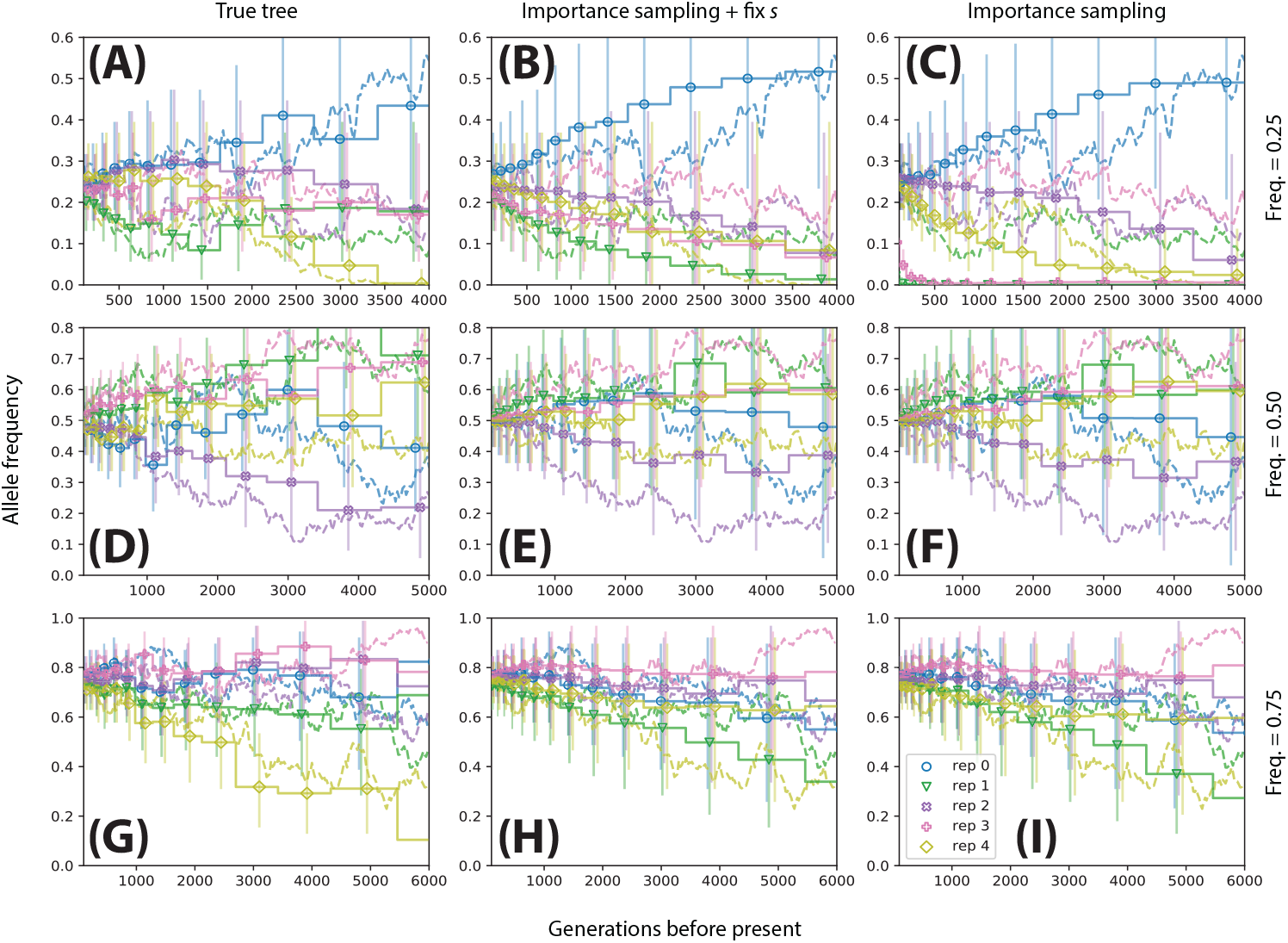
Performance of trajectory inference across replicates. We illustrated inferred vs. true trajectories, holding selection coefficient constant across the replicates. In this case, we chose *s* = 0 in order to maximize the amount of variability in the trajectories of the replicates. Rows correspond to simulations conditioned on different present-day allele frequencies (0.25: A-C,0.50: D-F,0.75: G-I). Left column (A,D,G): trajectories inferred using the true local tree. Middle column (B,E,H): trajectories inferred using importance sampling, fixing the selection coefficient to the ground truth (*s* = 0). Right column (C,F,I): trajectories inferred under importance sampling and estimating *s*. Simulations were done under the constant size model described in Methods and Materials using a locus of 100kb, *n* = 25 diploid individuals and *µ* = 2.5 × 10^*−*8^ mut/bp/gen, *r* = 1.25 × 10^*−*8^ recombinations/bp/gen.

**S7 Fig.**
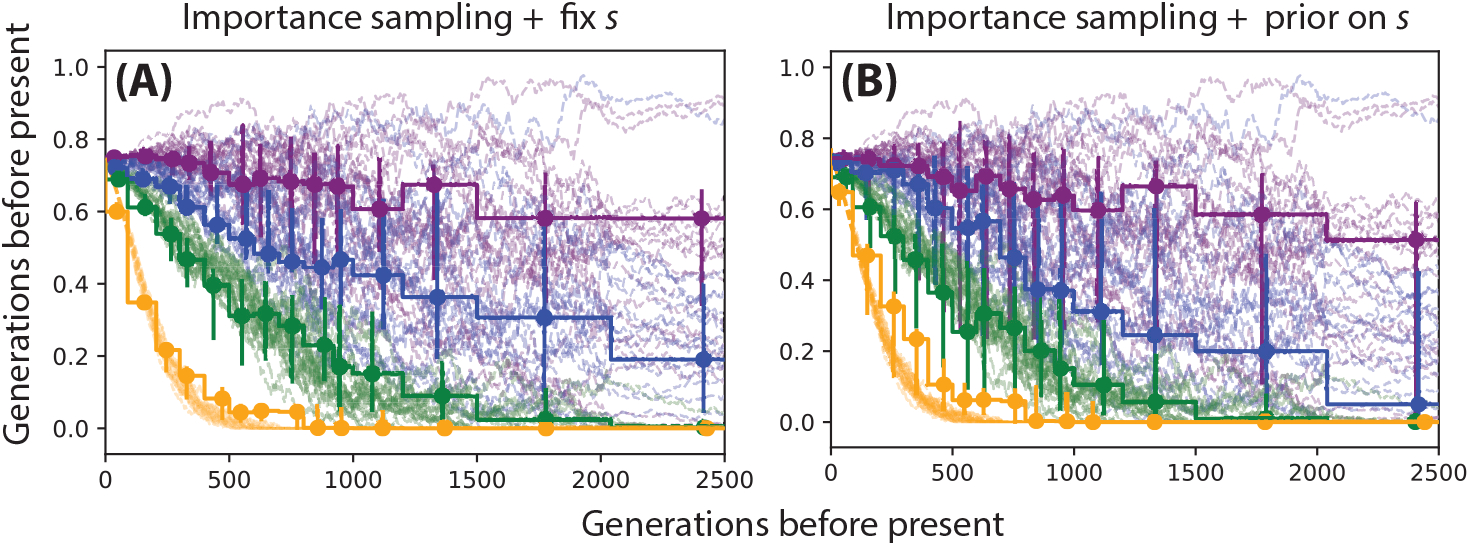
Effect of uncertainty in *s* on trajectory inference. A: trajectories inferred using importance sampling (i.e. Eq. 23), fixing *ŝ* = *s*. B: trajectories inferred using importance sampling and integrating over a uniform prior on *s* (see S1 Text for formulae). Simulations were done under the European demographic model described in Methods and Materials using a locus of 200kb, *n* = 25 diploid individuals and *µ* = 2.5 × 10^*−*8^ mut/bp/gen, *r* = 1.25 × 10^*−*8^ recombinations/bp/gen.

**S8 Fig.**
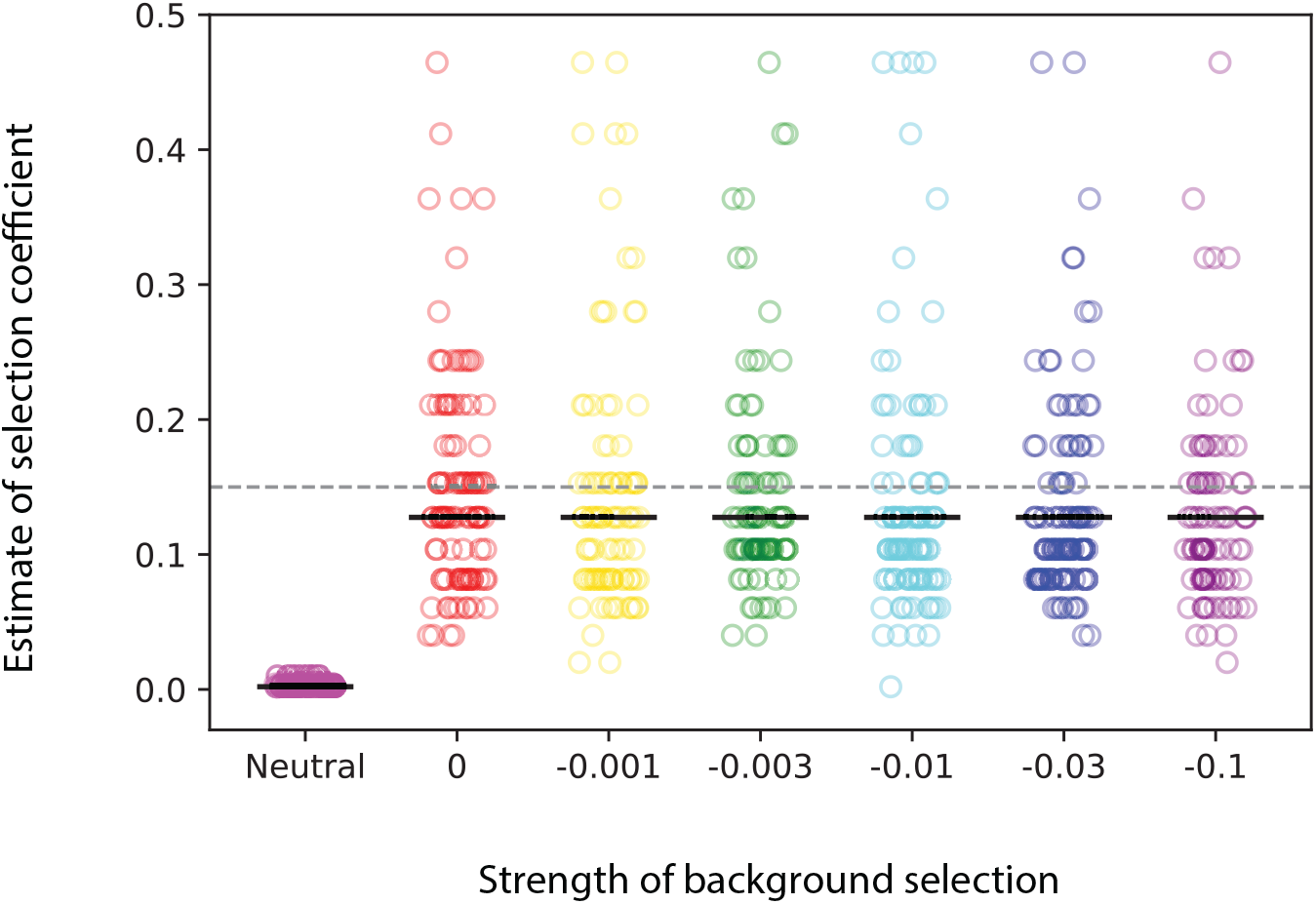
Effect of background selection on inference of selection. Estimates of the selection coefficient of a selected allele (*s* = 0.1) linked to sites under background selection of varying strength. Simulations were done using forward simulations assuming constant *N*_*e*_ = 10^3^ using a locus of 1Mb, *n* = 25 diploid individuals and *µ* = 2.5 × 10^*−*8^mut/bp/gen, *r* = 1.25 × 10^*−*8^ recombinations/bp/gen. Mean estimates are represented by blue hash marks.

**S9 Fig.**
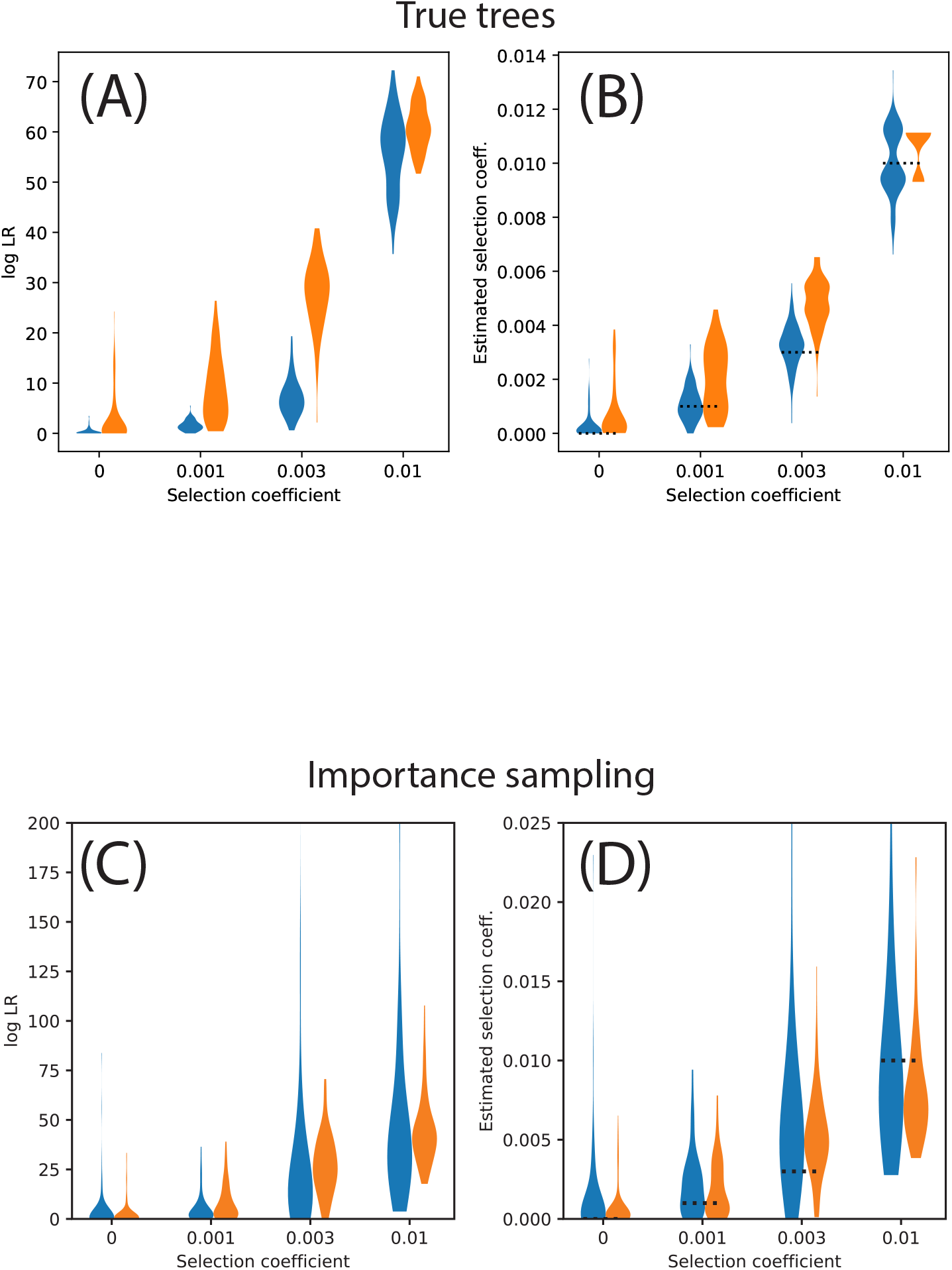
Effect of model misspecification on inference of selection. Likelihood ratios (A,C) and selection coefficient estimates (B,D) calculated given the true tree (A,B) and via importance sampling using ARGweaver (C,D). Blue violin plots represent estimates using the correct (European) demographic model, whereas orange plots represent estimates using a model of constant *N*_*e*_ = 10^4^. Simulations were done under the European demographic model described in Methods and Materials using a locus of 200kb, *n* = 25 diploid individuals and *µ* = 2.5 × 10^*−*8^ mut/bp/gen, *r* = 1.25 × 10^*−*8^ recombinations/bp/gen.

**S10 Fig.**
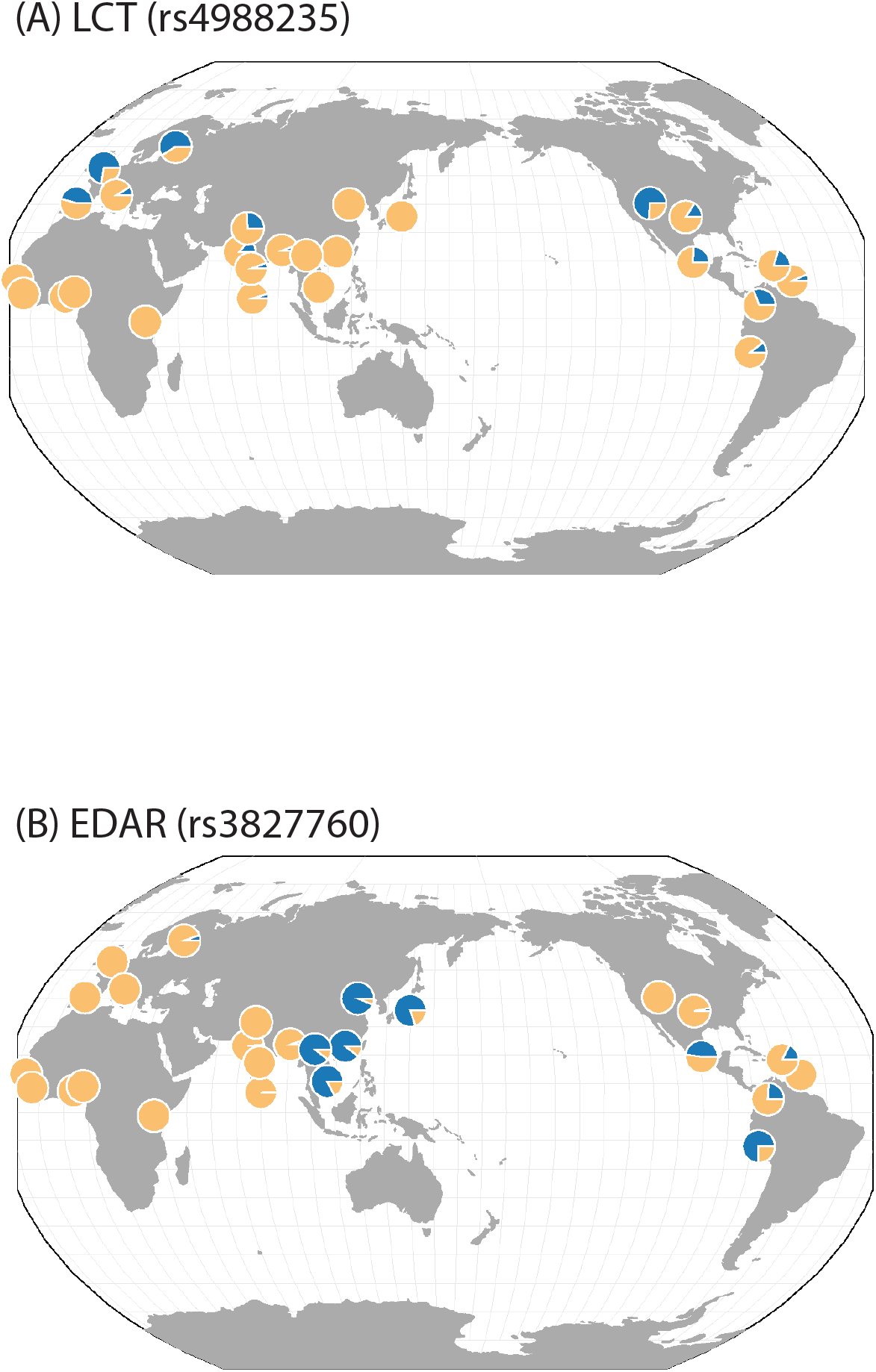
Geographical distribution of pigmentation SNPs. Population-wide allele frequencies of (A) rs4988235 (MCM6) and (B) rs3827760 (EDAR) plotted geographically using GGV.

**S11 Fig.**
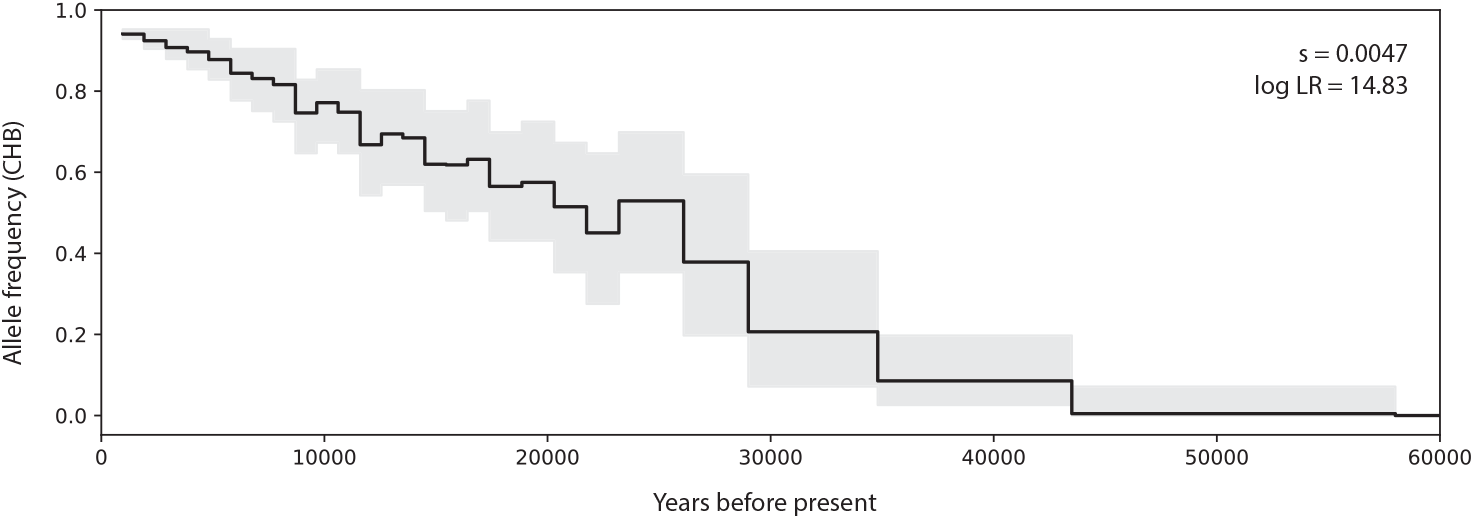
Allele frequency trajectory of rs3827760 (EDAR) in CHB. Frequency trajectory inferred using importance sampling under a model of CHB demography.

**S12 Fig.**
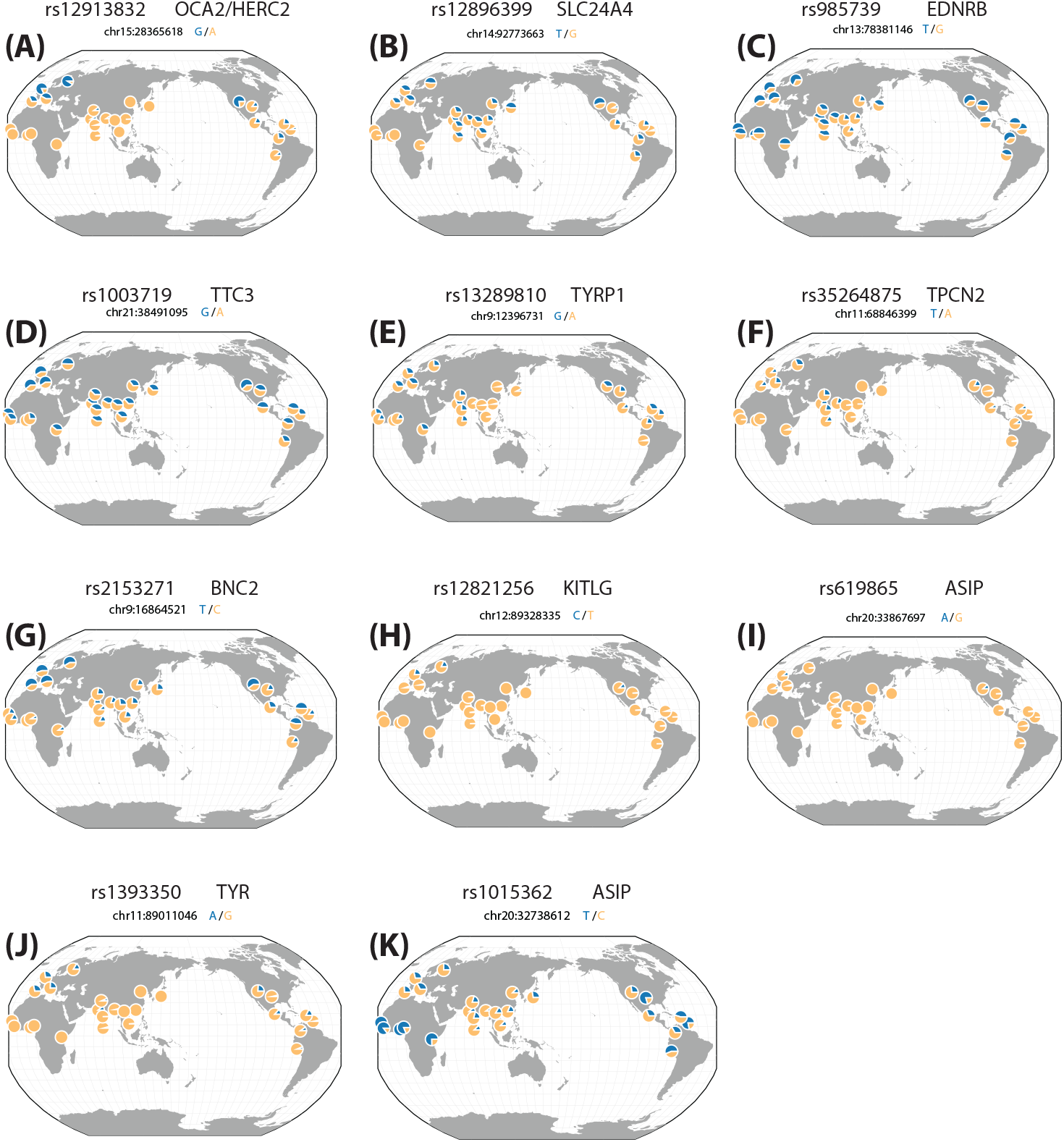
Geographical distribution of pigmentation SNPs. Population-wide allele frequencies of pigmentation SNPs from Fig. 10 plotted geographically using GGV.

**S13 Fig.**
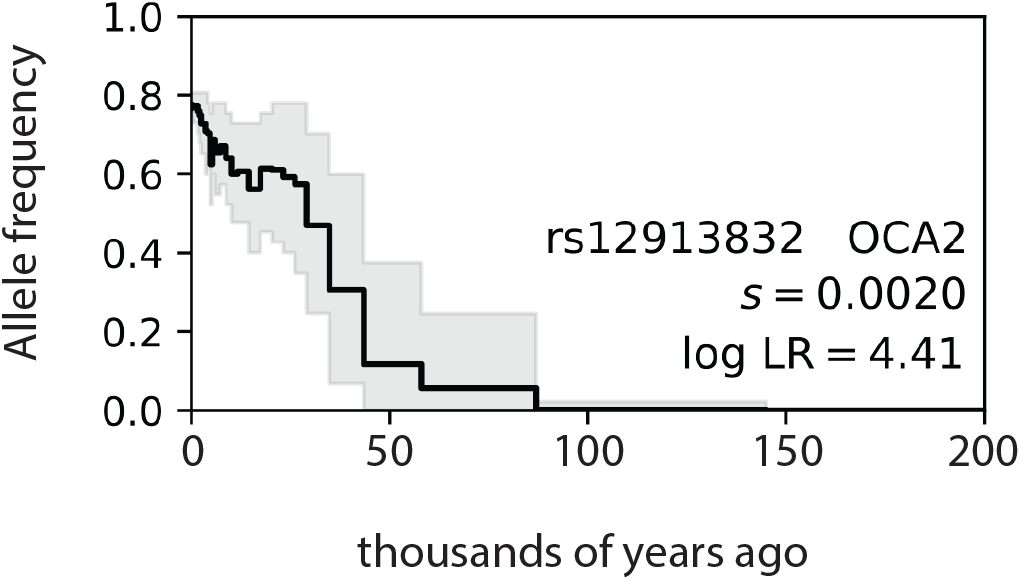
Allele frequency trajectory estimate of rs12913832 (OCA2/HERC2). The same trajectory estimate as in Fig. 10A with *x*-axis limits extended to illustrate earlier history of the allele.

**S1 Text. Appendix and commands to reproduce simulations and analyses.**

