## Supplementary material for "An approximate full-likelihood method for inferring selection and allele frequency trajectories from DNA sequence data": S1 Text

### Supplementary Text

Aaron J. Stern

July 22, 2019

### 1 Appendix

#### 1.1 Glossary of mathematical symbols

| symbol | domain | description |
|---|---|---|
| $s$ | $\mathbb{R}_{\geq 0}$ | the selection coefficient* |
| $K$ | $\mathbb{N}_{\geq 2}$ | the number of discrete timepoints |
| $X$ | $[0,1]^K$ | allele frequency trajectory |
| $N$ | $\mathbb{R}^K_{>0}$ | population size trajectory |
| $G$ | $\cdot$ | the ancestral recombination graph (ARG) |
| $G_k$ | $\cdot$ | the local tree at the site indexed by $k$ |
| $G_{\setminus k}$ | $\cdot$ | the ARG, omitting $G_k$ |
| $C^{\text{der}}$ | $\mathbb{N}^K$ | the number of derived lineages remaining at each timepoint $1,\ldots,K$. |
| $C^{\text{anc}}$ | $\mathbb{N}^K$ | the number of ancestral lineages remaining at each timepoint. |
| $C^{\text{mix}}$ | $\mathbb{N}^K$ | the number of mixed lineages remaining at each timepoint. |
| $C$ | $\cdot$ | $:= (C^{\text{der}}, C^{\text{anc}}, C^{\text{mix}})$ |
| $L(s)$ | $\mathbb{R}_{\geq 0}$ | the full likelihood of the selection coefficient $s$ |
| $M$ | $\mathbb{N}$ | the number of posterior ARG samples, after thinning and burn-in |
| $G^{(m)}$ | $\cdot$ | the $m$th posterior ARG sample, after thinning and burn-in s.t. $m \in 1,\ldots,M$ |
| $\widehat{\text{LR}}(s)$ | $\mathbb{R}_{\geq 0}$ | importance sampling estimate of $L(s)/L(s=0)$ |
| $\Omega^{(m)}$ | $\mathbb{R}_{\geq 0}$ | $m$th importance sampling weight |

*NB: as we mention in the main text, we also use $s$ as shorthand for arbitrarily complex parameters describing the selection model; e.g., $s$ (in the Methods derivations) can be thought of as shorthand for a vector containing both the actual selection coefficient as well as the timing of the onset of selection.

#### 1.2 Calculating allele frequency transition probabilities

Our likelihood calculations require allele frequency transition distributions for different selection coefficients, population sizes, and spans of time. Rather than employ the more common approach of numerically calculating allele frequency transition distributions using the Wright-Fisher diffusion process with drift and selection (e.g., [1,2]), we follow [3] and precompute allele frequency transition distributions on a grid of time spans (i.e., generations) and scaled selection coefficients (i.e., $\alpha = 2Ns$) using the Wright-Fisher model of reproduction in a finite population experiencing genetic drift and natural selection (see [1]). Specifically, for each value of $\alpha$, we use simple matrix multiplication to produce allele frequency transition matrices for discrete frequencies in a haploid population of size $N^{hap} = 2000$ at a number of generations spanning from $g = 1$ to $g = g_{\max}$ (corresponding to scaled drift times of 1/2000 to $g' = g_{\max}/2000$). We use this smaller population size to approximate the transition probabilities in larger population sizes ($N \geq 10^4$), where calculating the full Wright-Fisher transition matrix is prohibitively expensive. We approximate the probability of a transition over $t$ generations with selection coefficient $s$ under diploid population size $N$ using the haploid population of size $N^{hap}$ with rescaled time $\tilde{s} = \frac{N^{hap}}{2N}t$ and rescaled selection coefficient $\tilde{s} = \frac{2N}{N^{hap}}s$. Our simulated results suggest this model is accurately even when $\frac{2N}{N^{hap}} \approx 100$ (S3 Fig).

The allele frequency $X$ in the haploid population take on discrete values in $\{0, 1/N^{hap}, 2/N^{hap}, \ldots, 1\}$ Let $X_k$ be the allele frequency in the $k$th epoch. Then, conditional on $X_k = x_k$, $Y_{k+1} := N^{hap}X_{k+1}$ follows a binomal distribution $\text{Bin}\big(N^{hap}, p^{\ddagger}(x_k)\big)$, where

$$p^{\ddagger}(x) := p^{\dagger}(x)(1+s)/(p^{\dagger}(x)(1+s) + 1 - p^{\dagger})$$

and

$$p^{\dagger}(x) := (1-u)x + v(1-x)$$

and $u$ and $v$ are the mutation rates from derived to the ancestral type and vise versa, respectively. We note that $u$ and $v$ are also rescaled similarly to $s$ in order to approximate mutation in a population of smaller size. Thus, the transition probability from $i \rightarrow j$ is simply the probability $\text{Bin}\big(N^{hap}, p^{\ddagger}(x_k)\big)$.

The spacing of time points for these transition probabilities is chosen *a priori*; in practice, we use linear spacing for recent history and/or periods of population growth. We bin allele frequencies into $d$ discrete frequency categories unevenly distributed between 0 and 1 such that extreme frequency bins outnumber intermediate frequency bins. To calculate allele frequency transition distributions for time spans and selection coefficients not contained in the grid of pre-computed values, we linearly interpolate between the nearest precomputed values. See [3] for details.

We also note that if the time of the onset of selection, $t_s$, is to be inferred, then it is necessary to let $s$ depend on the epoch $i$; specifically, whether the allele is under selection vs. neutral during said epoch. Let $s_i$ denote the value of the selection coefficient during epoch $i$, and $\mathbf{s} = (s_1, \ldots, s_K)$.

Additionally, we condition the allele frequency process on the present-day frequency $X_0$ by using the following reweighting:

$$\mathbb{P}(X_i \mid X_{i+1}, X_0, \mathbf{s}) = \frac{\mathbb{P}(X_i \mid X_{i+1}, \mathbf{s})\mathbb{P}(X_0 \mid X_i, s)}{\mathbb{P}(X_0 \mid X_{i+1}, \mathbf{s}}$$

where $\mathbb{P}(X_{i_1} \mid X_{i_2}, s)$ is the forward-time unconditional probability of transitioning from $X_{i_2}$ to $X_{i_1}$ (in coalescent time, $t_{i_2} > t_{i_1}$; in forward time, $t_{i_2} < t_{i_1}$). Please see **??** for practical documentation on running this procedure.

#### 1.3 Forward and backward probabilities

Here we derive recursions for the forward and backward probabilities $f_i(x_i)$ and $b_i(x_i)$, respectively. These quantities are equivalent to $\mathbb{P}(C_{1:i} \mid X_i, N_{i-1})$ and $\mathbb{P}(C_{i+1:K-1}, X_i \mid X_i, N_i)$, respectively, where $C_{a:b} = C_a, C_{a+1}, \ldots, C_b$.

Let $b_i(x_i) = \mathbb{P}(C_{1:i} \mid X_i, N_{i-1})$. We calculate this quantity recursively moving from $i = 1 \to i$:

$$b_1(x_1) = \sum_{x_0} \mathbb{P}(C_1 \mid C_0, X_0 = x_0, N_0)\, \mathbb{P}(X_0 = x_0 \mid X_1 = x_1, N_0, s) \tag{1}$$

$$b_i(x_i) = \sum_{x_{i-1}} b_{i-1}(x_{i-1})\, \mathbb{P}(C_i \mid C_{i-1}, X_{i-1} = x_{i-1}, N_{i-1})\, \mathbb{P}(X_{i-1} = x_{i-1} \mid X_i = x_i, N_i, s) \tag{2}$$

and we can apply this recursion to calculate the likelihood function of $s$ given $G$ as

$$\mathrm{L}(s \mid G) \propto b_K(0). \tag{3}$$

The above is commonly known as the backward algorithm when applied to HMMs. In our model, the backward algorithm's recursion proceeds backwards through time. Alternatively, using the forward algorithm, with its recursion proceeding forwards in time:

$$f_{K-1}(x_{K-1}) = \mathbb{P}(X_{K-1} = x_{K-1} \mid X_K = 0, N_{K-1}, s) \tag{4}$$

$$f_i(x_i) = \mathbb{P}(C_{i+1} \mid C_i, X_i = x_i, N_{i+1}) \sum_{x_{i+1}} f_{i+1}(x_{i+1}) \mathbb{P}(X_i = x_i \mid X_{i+1} = x_{i+1}, N_i, s) \tag{5}$$

and we can apply this recursion to calculate the likelihood function of $s$ given $G$ as

$$\mathrm{L}(s \mid G) \propto \sum_{x_0} f_0(x_0) \tag{6}$$

#### 1.4 Importance sampling estimate of the posterior probability of the allele frequency

Here we show a quick derivation of the importance sampling estimate of the marginal posterior probability of the allele frequency trajectory at timepoint $i$, i.e. the posterior of $X_i$. Notation follows directly from the glossary.

First, let us establish that $\mathbb{P}(X_i \mid G_k, s) = \mathbb{P}(X_i \mid C_k, s)$; i.e., that the topology of the tree, conditioned on the allelic labeling of its leaves, does not affect the posterior probability of $X_i$. We will suppress $s$ for easy of notation.

$$\mathbb{P}(X_i \mid G_k) = \frac{\mathbb{P}(G_k \mid X_i)\mathbb{P}(X_i)}{\mathbb{P}(G_k)} \tag{7}$$

$$= \frac{\mathbb{P}(\text{topo}_k \mid C_k, X_i)}{\mathbb{P}(\text{topo}_k \mid C_k)} \frac{\mathbb{P}(C_k \mid X_i)\mathbb{P}(X_i)}{\mathbb{P}(C_k)} \tag{8}$$

Because the topology is independent of the allele frequency if we condition on the allelic labeling,

$$= \frac{\mathbb{P}(\text{topo}_k \mid C_k)}{\mathbb{P}(\text{topo}_k \mid C_k)} \frac{\mathbb{P}(C_k \mid X_i)\mathbb{P}(X_i)}{\mathbb{P}(C_k)} \tag{9}$$

$$= \frac{\mathbb{P}(C_k \mid X_i)\mathbb{P}(X_i)}{\mathbb{P}(C_k)} \tag{10}$$

$$= \mathbb{P}(X_i \mid C_k) \tag{11}$$

where we use topo to denote the topology of the tree, conditioned on its allelic labeling.

Next, we derive the importance sampling estimator of the allele frequency marginal posterior:

$$\begin{aligned}
\pi(X_i \mid D, s) &= \mathbb{E}_{G|D,s}[\mathbb{P}(X_i \mid G, D, s)] && (12)\\
&= \mathbb{E}_{G|D,s=0}\left[\mathbb{P}(X_i \mid G, D, s)\frac{\mathbb{P}(G \mid D, s)}{\mathbb{P}(G \mid D, s=0)}\right] && (13)\\
&= \mathbb{E}_{G|D,s=0}\left[\mathbb{P}(X_i \mid G, D, s)\frac{\mathbb{P}(G \mid s)}{\mathbb{P}(G \mid s=0)}\right] \times \frac{L(s)}{L(s=0)} && (14)\\
&\propto \mathbb{E}_{G|D,s=0}\left[\mathbb{P}(X_i \mid G, D, s)\frac{\mathbb{P}(G \mid s)}{\mathbb{P}(G \mid s=0)}\right] && (15)\\
&\approx \mathbb{E}_{G|D,s=0}\left[\mathbb{P}(X_i \mid G_k, G_{\setminus k}, D, s)\frac{\mathbb{P}(G_k \mid s)}{\mathbb{P}(G_k \mid s=0)}\right] && (16)\\
&\approx \mathbb{E}_{G|D,s=0}\left[\mathbb{P}(X_i \mid G_k, s)\frac{\mathbb{P}(G_k \mid s)}{\mathbb{P}(G_k \mid s=0)}\right] && (17)\\
&= \mathbb{E}_{G|D,s=0}\left[\mathbb{P}(X_i \mid C_k, s)\frac{\mathbb{P}(C_k \mid s)}{\mathbb{P}(C_k \mid s=0)}\right] && (18)
\end{aligned}$$

Hence,

$$\frac{1}{M}\sum_{m=1}^{M}\mathbb{P}(X_i \mid C_k^{(m)}, s)\Omega^{(m)}(s) \to \mathbb{E}_{G|D,s=0}\left[\mathbb{P}(X_i \mid C_k, s)\frac{\mathbb{P}(C_k \mid s)}{\mathbb{P}(C_k \mid s=0)}\right] \approx \kappa\pi(X_i \mid D, s) \qquad (19)$$

where $\kappa$ is $\left[L(s)/L(s=0)\right]^{-1}$, for which we have already established an importance sampling estimator (main text). Thus, our importance sampling estimate of the posterior marginal given $s$ is

$$\hat{\pi}(x_i \mid D, s) := \frac{\sum_{m=1}^{M}\mathbb{P}(X_i \mid C_k^{(m)}, s)\Omega^{(m)}(s)}{\sum_{m=1}^{M}\Omega^{(m)}(s)}. \qquad (20)$$

#### 1.5 Bayesian estimates of the selection coefficient

Allowing a prior distribution on $s$, $\pi(s)$, the posterior of the selection coefficient $\pi(s \mid D)$ follows

$$\pi(s \mid D) \propto \frac{L(s)}{L(s=0)}\pi(s) \approx \widehat{\mathrm{LR}}(s)\,\pi(s). \tag{21}$$

Then the estimate of the posterior marginal is given by

$$\hat{\pi}(x_i \mid D) = \int_{-\infty}^{\infty} \hat{\pi}(x_i \mid D, s)\pi(s|D)ds \tag{22}$$

which can be approximated by a sum over $d$ discretized values of $s$, $\mathcal{S} = \{s_1, \ldots, s_d\}$ as

$$\hat{\pi}(x_i \mid D) := \sum_{s \in \mathcal{S}} \hat{\pi}(x_i \mid D, s)\tilde{\pi}(s|D) \tag{23}$$

where $\tilde{\pi}$ represents a probability mass function over $s$.

### 2 Commands to reproduce simulations and analyses

#### 2.1 Simulations of trajectories, local trees, and haplotypes

To simulate data, we used a slight modification of the standard `discoal` package [4], available at `https://github.com/kern-lab/discoal`. In the standard `discoal` , there is no option to output the allele frequency trajectory, and there is also no option to simultaneously output the sample's local trees and haplotypes. This is important in order to compare inference vs. ground truth for the same replicate. Our modified version prints the trajectory to stdout, as well as the local trees and the haplotypes, and is available on the `CLUES` Github page. Nonetheless, the standard `discoal` documentation applies completely to our modified version, and we will leave the reader to learn the exact meaning of the arguments and options from documentation available through that repository.

To simulate data under the constant effective population size model, we ran

```
$ ./discoal 51 100 100000 -t 100 -r 50 -A 1 0 0.5 -x 0.5 -c 75e-2 -ws 0 -a 200 -N 10000 -
    i 4 > example.const.discoal
```

This specifies a sample of 50 modern haplotypes, simulated independently 100 times, with $N = 10^4$ diploid individuals, $4N\mu = 100$, $4Nr = 50$, and 1 ancient haplotype from 0.5 coalescent units ago. We specify the selected site to be in the center of the locus, segregating at 75% frequency in the present day, with a selection strength of $\alpha = 200 = 2Ns$ where $s = 0.01$. We simulate the trajectory assuming a time discretization of $1/(4N)$ coalescent units, on the order of 1 generation.

To simulate data under the European demographic model, we ran

```
$ ./discoal 51 100 100000 -t 3760 -r 1880 -A 1 0 0.021 -x 0.5 -c 75e-2 -ws 0 -i 4 -a 3762
    -N 188088 -en 0.000120 0 0.124319 -en 0.000272 0 0.042569 -en 0.000399 0 0.031529 -
    en 0.000532 0 0.023182 -en 0.000665 0 0.017045 -en 0.000797 0 0.012532 -en 0.000930 0
    0.009214 -en 0.001063 0 0.006576 -en 0.001224 0 0.009894 -en 0.001329 0 0.009894 -en
    0.001595 0 0.009894 -en 0.001994 0 0.009910 -en 0.002713 0 0.076953 -en 0.003722 0
    0.076953 -en 0.004918 0 0.076953 -en 0.006247 0 0.076892 -en 0.007870 0 0.038865 -en
    0.008507 0 0.038865 -en 0.009304 0 0.038865 -en 0.010367 0 0.038865 -en 0.011962 0
    0.038865 -en 0.014621 0 0.038865 -en 0.018608 0 0.038865 -en 0.023925 0 0.038865 -en
    0.033229 0 0.038865 -en 0.046521 0 0.038865 -en 0.066458 0 0.038865 -en 0.132917 0
    0.038865 -en 0.398750 0 0.038865 > example.ceu.discoal
```

This specifies a sample of 50 modern haplotypes, simulated independently 100 times, with $N =$ 188088 diploid individuals, $4N\mu = 3760$, $4Nr = 1880$, and 1 ancient haplotype from 0.021 coalescent units ago (scaled by the present-day effect population size, $N = 188088$). We specify the selected site to be in the center of the locus, segregating at 75% frequency in the present day, with a selection strength of $\alpha = 3762 = 2Ns$ where $s = 0.01$. We simulate the trajectory assuming a time discretization of $1/(4N)$ coalescent units, on the order of 1 generation. We use the `-en` option in order to scale effective population size to the harmonic mean of the population size during that time interval.

```
$ ./discoal 101 100 100000 -t 3760 -r 1880 -A 1 0 0.021 -x 0.5 -c 50e-2 -ws 0 -a 3762 -f
    0.268 -i 4 -N 188088 -en 0.000120 0 0.124319 -en 0.000272 0 0.042569 -en 0.000399 0
    0.031529 -en 0.000532 0 0.023182 -en 0.000665 0 0.017045 -en 0.000797 0 0.012532 -en
    0.000930 0 0.009214 -en 0.001063 0 0.006576 -en 0.001224 0 0.009894 -en 0.001329 0
    0.009894 -en 0.001595 0 0.009894 -en 0.001994 0 0.009910 -en 0.002713 0 0.076953 -en
    0.003722 0 0.076953 -en 0.004918 0 0.076953 -en 0.006247 0 0.076892 -en 0.007870 0
    0.038865 -en 0.008507 0 0.038865 -en 0.009304 0 0.038865 -en 0.010367 0 0.038865 -en
    0.011962 0 0.038865 -en 0.014621 0 0.038865 -en 0.018608 0 0.038865 -en 0.023925 0
    0.038865 -en 0.033229 0 0.038865 -en 0.046521 0 0.038865 -en 0.066458 0 0.038865 -en
    0.132917 0 0.038865 -en 0.398750 0 0.038865
```

This specifies the same demographic model as in the previous simulation, except we increase the sample size to 100 haplotypes (and still 1 ancient haplotype). Additionally, to enforce a SSV, we use `-f 0.268` to enforce that the allele evolves under selection from the present day back to the point that it reaches a frequency of 0.268, and neutrally leading up to that point. We must simulate the SSV this way because `discoal` does not have an option to specify the time of selection's onset. We obtained the frequencies for the `-f` option by simulating under selection and finding the average frequency of the allele 100 generations before the present.

#### 2.2 Reformatting `discoal` output

It is necessary to parse the output of `discoal` to not only prepare the input files for `ARGweaver` (and `CLUES` , which just uses `ARGweaver` -formatted data), but also useful to separate trajectories, local trees, and haplotypes into separate data files. We wrote a Python script to run this process, `parseDiscoalOutput.py`, which is available on our Github page. The command to run this script is

```
$ python parseDiscoalOutput.py example.discoal <length_of_sequence> <num_sites> <num_haps
    > <out>
```

where you set the arguments to be the length of the sequence in base-pairs, the number of sites in the `discoal` simulation (in the two examples above, this would be $10^5$), the total number of haplotypes sampled ($n = 51$), and a basename for output files. This script will generate 3 files, with extensions `.traj`, `.trees`, `.sites`, that hold the trajectory, the true local tree at the site of interest, and the haplotypes reformatted in `ARGweaver` format, respectively. These files will be generated and named by the index of each replicate simulated in the file `example.discoal`. Note that currently this script is hardcoded to assume the SNP of interest is located at the center of the locus.

#### 2.3 Performing ARG-sampling using `ARGweaver`

We use the `arg-sample` function in the `ARGweaver` package, available at `https://github.com/mjhubisz/argweaver`, to sample the posterior ARG [5]. This function requires one major input: the `.sites` we generated in the previous step. However, it is also necessary to provide the proper demographic model, mutation rate, and recombination rate. Furthermore, you should specify the desired length of the MCMC chain (here $M = 3000$ samples). You can also compress sequence blocks to greatly speed up the process (here we compress down to 25-bp blocks). By default, ARGweaver thins down to every 10th sample, but this option may be adjusted.

To sample ARGs under a constant population size, we run

```
$ ./arg-sample -s example.const.sites -o example.const --age-file N_10000_agefile.txt --
    times-file N_10000_timesfile.txt -N 10000 --overwrite --quiet -m 2.5e-8 -r 1.25e-8 -c
     25 -n 3000 --resample-window 40000 --resample-window-iters 8 --infsites
```

To sample ARGs under the European demographic model, we run

```
$ ./arg-sample -s example.ceu.sites -o example.ceu --times-file tennessen_times_fine.txt
    --age-file tennessen_age.txt --popsize-file tennessen_popsize_fine.txt --overwrite -m
     2.5e-8 -r 1.25e-8 -c 25 --quiet -n 3000 --resample-window 100000 --resample-window-
    iters 8 --infsites
```

The files that are specified using `--times-file`, `--age-file`, and `--popsize-file` correspond to specifying the time discretization (in generations), the age of the ancient haplotype used to polarize alleles (in generations), and the population size trajectory. We supply all of the corresponding files on the `CLUES` Github page.

We also want to point out several tuning parameters: `--resample-window` and `--resample-window-iters` adjust the size of the resampling window and the number of resamples to perform on a particular window. Adjusting these parameters can affect the behavior of the MCMC routine by changing how aggressively changes are proposed to the ARG. Increasing the resample window will decrease the acceptance probability of a given proposal, but increasing the number of iterations will increase that probability that any of these proposals will be accepted. These parameters should be adjusted to yield about a 30-70% acceptance rate (Melissa Hubisz, personal communication).

This procedure will output a series of `.smc.gz` files.

#### 2.4 Extracting local trees from `ARGweaver` samples

We used the `smc2bed-all` and `arg-summarize` programs included in the `ARGweaver` package to extract local trees at the site of selection. Your `ARGweaver` output has the form `example.<k>.smc.gz`, where $\texttt{k} = 0, 10, 20, \ldots, 3000$. To run extraction,

```
$ ./smc2bed-all example; ./arg-summarize -a example.bed.gz -r chr:50000-50000 -l example.
    log -E > argweaver.example.trees
```

This saves a list of Newick trees extracted from the site 50000 to `argweaver.example.trees`.

#### 2.5 Preliminaries for `CLUES`

`CLUES` depends on a probabilistic model for allele frequency changes. Thus, it is necessary to either download our pre-computed transition probabilities for either the constant $N = 10^4$ or European demography models (formatted in HDF5 using the `h5py` package [6]). We provide an example file `example.f_75.hdf5`, precomputed conditioned on $X(0) = 0.75$, but one can alternatively compute transition probabilities from scratch for a custom model. We next describe how to do so.

To compute transition probabilities for a set of selection coefficients `s1,s2,...sL` from scratch, run the following commands:

```
$ python make_transition_matrices_from_argweaver.py <Nsmall> <s1> example.log trans.s_<s1
    >.h5 --breaks 0.95 0.025 --debug
```

```
$ python make_transition_matrices_from_argweaver.py <Nsmall> <s2> example.log trans.s_<s2>.h5 --breaks 0.95 0.025 --debug
...
$ python make_transition_matrices_from_argweaver.py <Nsmall> <sL> example.log trans.s_<sL>.h5 --breaks 0.95 0.025 --debug
$ mkdir example_trans_dir; mv trans.s_*.h5 example_trans_dir
```

The argument `Nsmall` denotes the population size of the Wright Fisher model used to calculate the times. It should be no greater than $\sim 10^4$, and only around $\sim 10^3$ if you want it to run quickly; note that this number can be mucher smaller than the "true" population size, and it is scaled down to simply speed up calculations, and results are rescaled to the "true" size. The `example.log` file is obtained from the `ARGweaver` run, and it summarizes the actual demographic model.

After completing this step, it is necessary to aggregate the transition probabilities and condition them on present-day frequencies:

```
$ python conditional_transition_matrices.py example.log example_trans_dir/ --listFreqs 0.25 0.5 0.75 -o trans
```

This will create a HDF5 file called `trans.hdf5`. This file contains transition matrices conditioned on the present-day frequency being 0.25, 0.50, and 0.75. It may be wise to use a richer set of frequencies by modifying `--listFreqs` if you are interested in analyzing real data.

To calculate transition matrices under an SSV model, there is a `--ssv` option. Warning: this will require substantially longer runtime and storage than the model assuming a hard sweep.

#### 2.6 Running `CLUES`

To run `CLUES` , the minimal command is

```
$ python clues.py <treesFile> <conditionalTrans> <sitesFile> <popFreq>
```

For example,

```
$ python clues.py example.trees example.f_75.hdf5 example.sites 75e-2 --thin 10 --burnin 100 --output example.clues
```

The 4 key inputs here are:

1. `treesFile`: local trees sampled and extracted from `ARGweaver` .

2. `conditionalTrans`: transition matrices conditioned on selection coefficient and present-day frequency, formatted in HDF5.

3. `sitesFile`: the `.sites` file used by `ARGweaver` . (This file is necessary to label the trees by derived/ancestral allele.)

4. `popFreq`: the present-day allele frequency you'd like to condition on. If `conditionalTrans` is conditioned on too sparse a grid of present-day frequencies, `CLUES` will fail if `popFreq` is too distant from all of the frequencies.

There are also several options you can deploy. Here we show `--thin` and `--burnin`, which we use here to treat the first 100 trees in `example.trees` as burnin, and thin down to every 10th tree in the file after that point. This corresponds to an overall burnin of 1000 samples and an overall thinning rate of 100 trees, assuming you use the baseline `ARGweaver` thinning rate of 10 trees. The `--output` option saves the output of `CLUES` (e.g. the likelihood surface, importance sampling weights, MLEs, trajectory posterior marginals, and more) to a HDF5 file named `example.clues.h5`.

There are more options available. To run `CLUES` under an SSV model, simply use the `--ssv` option. (Note: running `--ssv` will require a transition probability file computed using the `--ssv` option in `conditional_transition_matrices.py`.) To fix the value of $s = s'$ (rather than optimize over all potential values of $s$), use the option `--selCoeff s'`. To deploy a uniform prior on $s$, use the option `--prior`. To specify the position of the site of interest, set `--posn <position>`, which defaults to 50000.

#### 2.7 Runtime

To give a sense of the expected runtime of transition probability pre-computation, ARG sampling in `ARGweaver` , and `CLUES` itself, we timed each of these 3 steps for an example analysis. We found that transition probabilities ran in 36 minutes (14 minutes of unconditional transition probabilities, 22 minutes of conditioning; we used 45 discretized allele frequencies, 39 discrete timepoints, and 25 different values of the selection coefficient, on par with values used in our analysis in the main text). We ran ARG sampling and `CLUES` on the dataset used in the our study of background selection (see "Effects of background selection", main text). Time required to perform ARG sampling varied across replicates, but generally fell within 40-60 minutes. Time required to run `CLUES` was 5 minutes, using $M = 40$ sample ARGs after thinning. For analyses of larger regions and/or sample sizes, the ratio of `ARGweaver` runtime to `CLUES` runtime will increase.

6. Collette A. HDF5 for Python;. Available from: `https://www.h5py.org/`.
